# Redefining rock doves, *Columba livia*, using historical whole genome sequences

**DOI:** 10.1101/2023.04.12.536150

**Authors:** Germán Hernández-Alonso, Jazmín Ramos-Madrigal, George Pacheco, Hein van Grouw, Emily Louisa Cavill, Marta Maria Ciucani, Mikkel-Holger S. Sinding, Shyam Gopalakrishnan, M. Thomas P. Gilbert

## Abstract

The domestic pigeon’s exceptional phenotypic diversity was key in developing Darwin’s Theory of Evolution and establishing the concept of artificial selection in domestic species. However, unlike its domestic counterpart, its wild progenitor, the rock dove *Columba livia*, has received considerably less attention. Therefore, questions regarding their domestication, evolution, taxonomy, and conservation status remain unresolved. We generated whole-genome sequencing data from 65 historical rock dove samples representing all currently recognised subspecies and spanning the species’ original geographic distribution. Our dataset includes three specimens from Darwin’s collection and the type specimens of five different taxa. We characterised rock doves’ population structure, genomic diversity and gene-flow patterns. We show the West African subspecies *C. l. gymnocyclus* is basal to rock doves and domestic pigeons. Our results show gene-flow signals between the rock dove’s sister species *C. rupestris* and all rock doves except the West African populations. Our results led us to propose an evolutionary model for the rock dove considering the Pleistocene refugia theory. We propose that today’s rock dove genetic diversity and introgression patterns derive from a history of allopatric cycles and dispersion waves during the Quaternary glacial and interglacial periods. To explore the rock dove domestication history, we combined our new dataset with available genomes from domestic pigeons. Our results point to at least one domestication event in the Levant region that gave rise to all domestic breeds analysed in this study. Finally, we propose a species-level taxonomic arrangement to reflect the evolutionary history of the West African rock dove populations.

## Introduction

The pigeon *Columba livia*, is one the most common and familiar birds worldwide due to the ubiquitous distribution of its feral populations and the extended breeding practices of its domestic form. The domestic pigeon, on occasions bred to extreme exuberance (fancy breeds), has been an important model organism for the study of evolution, behaviour, genotype expression, among other research areas (Helms and Brugmann 2007). In contrast to the domestic pigeon, its parental species, the rock dove, has received relatively little attention and consequently, several unresolved questions remain regarding their taxonomy, evolution, conservation status, and domestication process.

Archeological evidence suggests that the rock dove was first domesticated between 10,000 to 3,000 years ago in either the Mediterranean Basin (Stringham et al. 2012; Johnston and Janiga 1995; Johnston 1992), or alongside the Neolithic Revolution in the Fertile Crescent (Driscoll et al. 2009). As opposed to deriving from a single domestication event, it has also been argued that rock doves were probably domesticated several times and in different places spanning their natural distribution range, possibly over a period of thousands of years (Shapiro and Domyan 2013; Johnston and Janiga 1995; Johnston 1992). After an early domestication phase, domestic pigeons were spread by humans throughout Eurasia and North Africa, where they started to locally diversify (Pacheco et al. 2020). Today, over 350 different domestic breeds are recognised, with the geographic origin for the major breed groups believed to be in the Middle East and South Asia. However, disentangling the rock dove’s domestication process is challenging, due to a history of extensive hybridisation and introgression both among domestic breeds, and between domestic and wild populations (Shapiro et al. 2013). Ultimately, there is a growing interest in improving our understanding of the rock dove’s evolutionary history, as such knowledge could improve their use as model organisms by understanding the origin of their genetic variation and the changes involved in their domestication process, or to contrast the effects of feralisation as a demographic and evolutionary process in the different feral pigeon populations.

Occurrence data suggest that the historical native distribution range of rock doves covered extensive areas of Europe, North Africa, the Middle East and South Asia, as well as sea coasts from the Faroe Islands and Britain, to Madeira, the Canaries, and the Cape Verde islands, mainly associated to nesting sites on rock faces and cliffs in coastal or montane regions (Urban et al. 2014; Shapiro and Domyan 2013; Gibbs et al. 2001; Johnston and Janiga 1995; Johnston 1992; Cramp 1985). However, given their long domestication history and the difficulties distinguishing wild from feral populations, it is challenging to accurately estimate their present or past distributions. For example, in some regions such as most of Continental Europe, the extirpation of natural populations, or the complete absorption into the gene pool of the expanding feral birds, considerably hinders the identification of the geographic range of wild rock doves (Gibbs et al. 2001; Johnston and Janiga 1995; Cramp 1985). Furthermore, it has even been suggested that true wild rock dove populations only remain in locations that are outside of the range of feral populations (Johnston and Janiga 1995; Johnston 1988). However, one recent genomic study on rock doves from the British Isles suggests this may not be the case, as all samples analysed from one of the regions believed to harbour wild populations contained genetic admixture from feral and/or domestic pigeons (Smith et al. 2022).

Phenotypic variability such as the size and colouration of the rock dove is geographically structured (Cramp 1985) and has been used to categorise their populations into subspecies (Johnston 1992). At present, most authors recognise nine subspecies of rock dove: *C. l. livia, C. l. gaddi, C. l. palestinae, C. l. schimperi, C. l. targia, C. l. dakhale, C. l. gymnocyclus, C. l. neglecta*, and *C. l. intermedia* (Goodwin 1977). Additionally, at least three more inconclusive subspecies have been proposed, *C. l. canariensis* from Canaries Islands, *C. l. atlantis* from Azores Islands, Madeira and Cape Verde, and *C. l. nigricans* from Mongolia and Northwest China (Urban et al. 2014; Gibbs et al. 2001; Cramp 1985). The debate about the last-mentioned subspecies revolves around their possible feral origins (Dickinson and Remsen 2013) or, in the case of the Subtropical Atlantic Islands rock doves, whether they are simply morphological variants of the *C. l. livia* subspecies (Gibbs et al. 2001; Cramp 1985; Murton and Clarke 196*8*).

Nevertheless, given that the morphological characteristics used to describe rock dove subspecies are clinally distributed, it is challenging to define the range limits of the subspecies, which are sometimes morphologically indistinguishable between adjacent populations (Goodwin 1977). Thus, there is little consensusabout the subspecies distribution ranges (Johnston 1992). Taking this into consideration, as well as the contiguously continental distribution of rock doves, the currently recognised subspecies could be questioned in terms of their evolutionary relevance (Mayr 1982).

A direct consequence of the limited knowledge, but also, the limited interest about the rock dove, is the lack of information related to their conservation status. Paraphrasing Johnston (1988), if we do not know how many rock doves exist at different places, we cannot be aware of the loss of populations; therefore, we cannot take actions to preserve them. The rock dove’s natural habitat is at risk due to human expansion, including remote areas where the rock dove could still survive (Johnston and Janiga 1995; Johnston 1992), but their main threat is the dissolution of their genetic pool resulting from genetic introgression from feral pigeons. These factors could lead to the imminent extinction of the rock dove within this century (Johnston and Janiga 1995; Johnston 1992; Johnston 1988).

To explore the rock dove diversity and their evolutionary history, we generated whole-genome sequencing data from 65 historical rock doves collected between 1865 and 1986, at localities spanning the species’ original distribution range. These samples represent all currently recognised subspecies. Our dataset includes 3 specimens from Darwin’s collection (collected in Madeira, Sierra Leone and the Shetland Islands), and the type specimens of 5 different taxa: *C. l. atlantis* Bannerman, 1931, *C. l. canariensis* Bannerman, 1914, *C. l. butleri* Meinertzhagen, 1921 (currently merged with *C. l. shimperi*), *C. l. dakhlae* Meinertzhagen, 1928, and *C. l. lividor* Bates, 1932 (currently merged with *C. l. gymnocyclus*). We used these genomes to study the diversity of rock dove populations, under the expectation of finding lower levels of admixture in historical (as opposed to modern) specimens, particularly in populations distributed far from human influence at that time. Simultaneously, we used our new dataset in combination with previously published genomes from domestic pigeons to re-investigate the origin of the domestic pigeon lineages. Moreover, we used our results to propose a model for the evolution of rock doves and their domestication. Finally, our results allowed us to confirm the feral origin of the subspecies *C. l. atlantis* and *C. l. canariensis*, identify the possible feral origin of *C. l. dakhlae* subspeices, and propose that *C. l. gymnocyclus* should be considered a full species, *Columba gymnocyclus* Gray, 1856.

## Results

### Rock dove genetic population structure follows geography

We produced whole genome sequencing data from 65 historical samples spanning the historical rock dove geographical range (Figure 1) to a depth of coverage of 0.8-7.89× (Table S1). To account for the low depth of coverage of some samples, a pseudo-haploid SNP panel was created using the historical samples together with genomic data from 39 publicly available domestic pigeons, 2 modern feral pigeons, 5 related Columbidae species (*Patagioneas fasciata, Columba palumbus, Columba larvata, Columba guinea, Columba rupestris*) and 1 common pheasant (*Phasianus colchicus*) (Table S2). To avoid the introduction of potential biases associated with historical DNA degradation, transitions were removed, resulting in a final dataset of 1,642,881 transversion sites.

**Figure 1.**
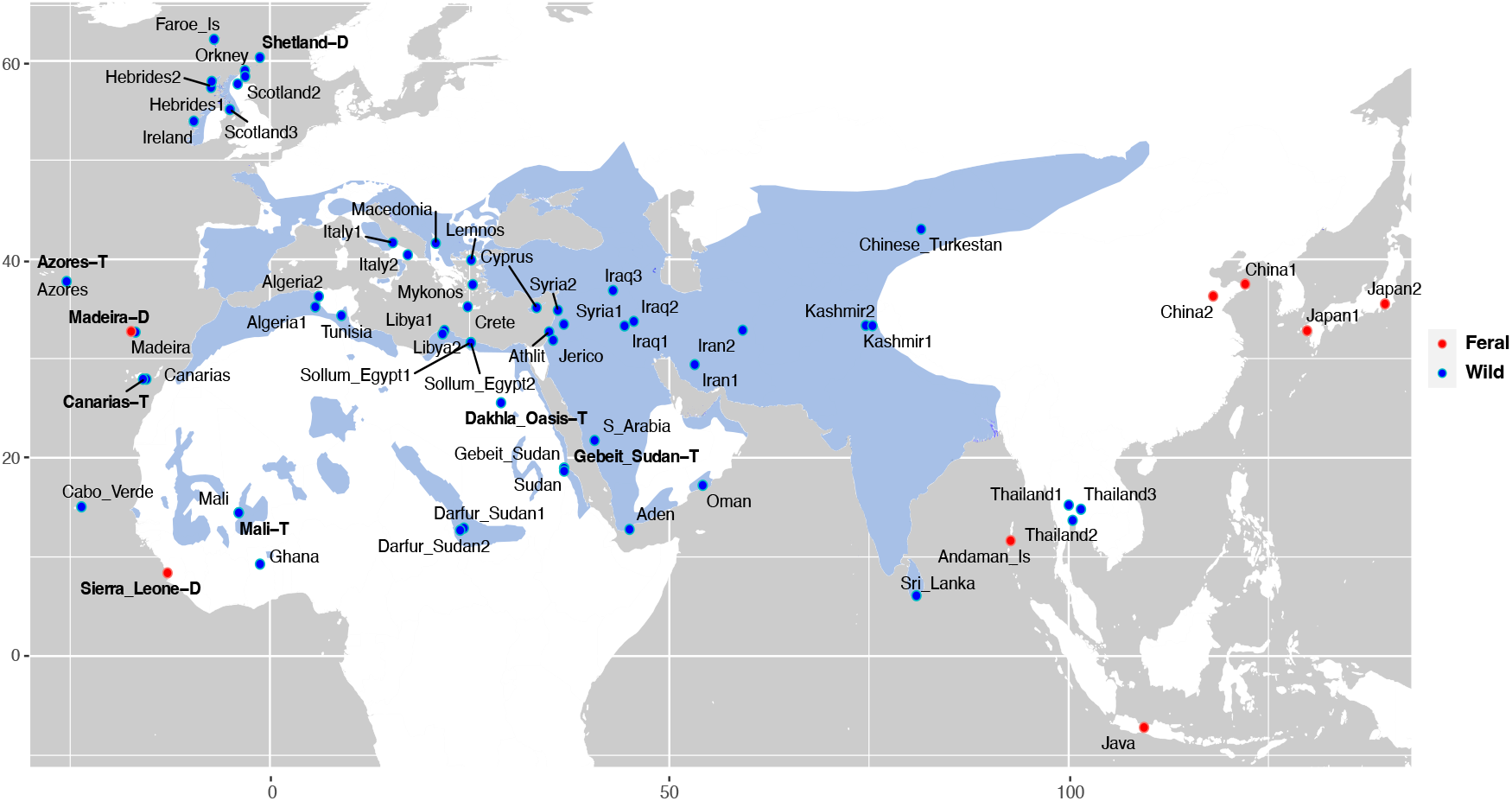
Geographic location of rock dove samples sequenced in this study. The map shows the approximate location of the wild historical rock doves and feral pigeons taking into consideration the annotated location associated with each museum specimen. The rock dove estimated area of occupancy according to The IUCN Red List of Threatened Species is depicted by the shaded blue area. Darwin’s collection specimens (D) and the type specimens (T) are shown in bold.

As an initial step to explore the genetic diversity of rock doves, we performed a multidimensional scaling analysis (MDS). Broadly, the majority of the samples are distributed across dimension 1, with the domestic pigeons at the right side of the plot and the historical rock doves at the left side. Several of our historical samples that were originally labeled in the collections as ‘feral’ are placed in the middle of the two clusters, as could be expected from admixed individuals. A divergent third cluster at the top of the plot that differentiates along dimension 2 includes the historical rock doves from the western part of Africa (Mali, Ghana), thus containing not only individuals of the *C. l. gymnocyclus* subspecies but also the type specimen for *C. l. lividor* (Mali-T) (Figure 2A).

**Figure 2.**
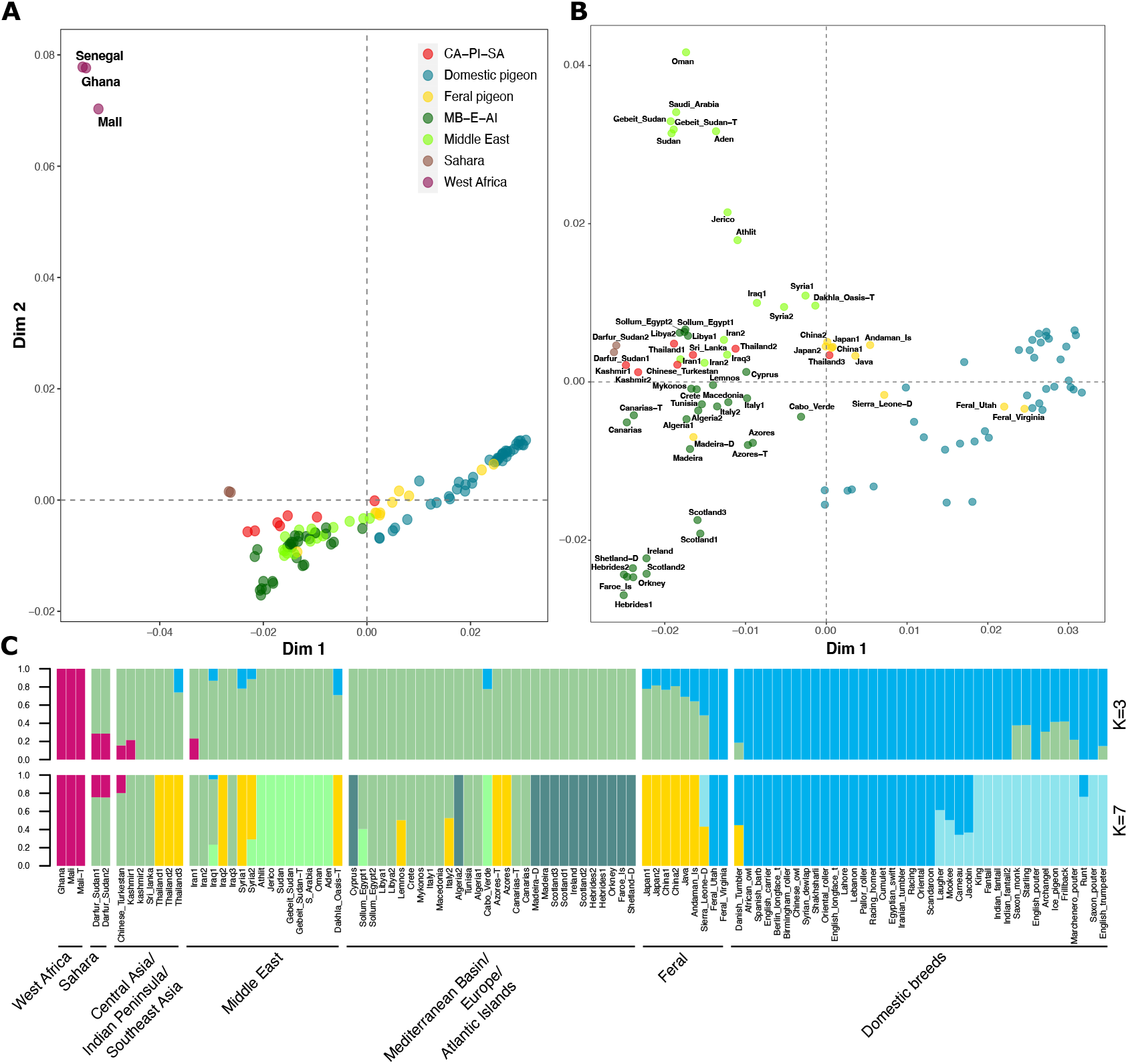
Rock dove’s population structure reveals a divergent population in West Africa. A-B) Multidimensional scaling (MDS) plots generated from pseudo-haploid data (1,642,881 transversion sites). A) MDS plot with all samples in our dataset, excluding the outgroups. Domestic breeds and historical rock doves are separated by dimension 1. West African rock doves appear divergent and separate on dimension 2. Colours represent the feral and domestic pigeons, as well as the geographic origin of the rock doves. Red colour includes samples from Central Asia, Peninsular India, and Southeast Asia (CA-PI-SA); dark green includes samples from the Mediterranean Basin, Europe, and the Atlantic Islands (MD-E-AI). B) MDS plot excluding the divergent West African rock doves, showing the population structure among the rock doves. C) Model-based clustering using ADMIXTURE assuming 3 and 7 ancestry components (K). Horizontal bars show different samples. Different colours show the estimated ancestry components and corresponding proportions. At K=7. Darwin’s collection specimens (D) and the type specimens (T) are shown in bold.

To explore the population structure of the rock dove and pigeon genomes in more detail, we estimated a second MDS plot excluding the West African rock doves (Figure 1B). Similar to the previous MDS, domestic pigeons and historical rock doves are separated in dimension 1, with the historical feral pigeons placed between these two clusters. We also observed that some of the historical rock doves that were originally described as wild based on their geographic location and morphological data, fall within the distribution of feral pigeons. This suggests they are in fact either feral, or admixed. For example, the *C. l. dakhlae* type specimen (Dakhla_Oasis-T) is placed close to the feral pigeons. The internal structure among the historical rock doves is defined by dimension 2, with the North Atlantic Islands rock doves (including the Shetland specimen from Darwin’s collection) in one end of the plot, and the rock doves from the Red Sea and Arabian Peninsula (including the type specimen of *C. l. butleri*, Gebeit_Sudan-T) in the other end. The remaining samples placed at the center of the historical rock dove cluster are mainly from the Mediterranean Basin, reflecting a geographical distribution along dimension 2 from northwest Africa (Algeria and Tunisia) going through the central Mediterranean, until the eastern Mediterranean and Near East. Subtropical Atlantic Islands rock doves are placed at the bottom part of this cluster, close to the northwestern African rock doves, and separated by dimension 1. Also placed in the Subtropical Atlantic Islands rock dove group, are the Darwin’s collection specimen Madeira-D labeled as feral, as well as the type specimens for the subspecies *C. l. canariensis* (Canarias-T) and *C. l. atlantis* (Azores-T) (Figure 2B).

To further explore the population structure in the dataset, we performed an admixture analysis to estimate individual ancestries assuming between 2 and 10 ancestral components (Figure 1C). When 2 ancestral components were estimated, we obtain one component that is mostly represented among domestic pigeons, and a second that is predominantly present among historical rock doves. Historical feral pigeons carry ancestry components from both clusters. In line with the MDS analysis, some of the historical rock doves (those placed close to the historical feral pigeons in the MDS plot) carry a fraction of ancestry from the same component identified in the domestic pigeons, including the *C. l. dakhlae* type specimen Dakhla_Oasis-T (Figure S3).When we extended the analysis to 3 ancestry components, all the West African rock doves (whether labeled as *C. l. gymnocyclus* or *C. l. lividor)* form a new independent cluster, consistent with the aforementioned results of the MDS analysis. Even though most samples retained similar ancestry proportions when assuming 2 or 3 components, some historical rock doves present a fraction of ancestry from the West African cluster, with the genomes of the rock doves from Darfur, in western Sudan, exhibiting the highest proportions (Figure 2C). Finally, when estimating 7 ancestry components, the specimens are split into clusters that resemble the geographical patterns found in the MDS plot (Figure 2B). The West African rock doves carry an ancestry component that is only shared with the samples from Darfur, Sudan (Figure S3). The feral birds get their own ancestry component, potentially representing the mixture of wild and domestic ancestry, with the samples from Thailand, Cape Verde, Azores (including the Azores-T type specimen), the Dakhla Oasis (Egypt), and one Iraqi sample (Iraq2) also carrying a proportion of this ancestry. Some other samples share this feral ancestry component, which could reflect different levels of domestic pigeon admixture on wild populations. The other ancestry components recovered among the historical rock doves broadly correspond to: 1) rock doves from the Near East and the Arabian Peninsula/Red Sea (including the Gebeit_Sudan-T type specimen); 2) rock doves from the North Atlantic Islands and Madeira (including the Darwin’s specimens from Shetland and Madeira, and the samples Cyprus and Algeria2), and; 3) rock doves from the Mediterranean Basin, Asia, and Canary islands (including the Canarias-T type specimen), as well as Darfur, Sudan (Figure 2C).

### Rock doves evolutionary relationships reveal the basal placement of the West African populations

To understand the evolutionary relationships among the historical rock doves, feral pigeons, and the domestic breeds, we built a Neighbor-Joining (NJ) tree based on identity-by-state pairwise distances (Figure 3A and S4). The internal branches of the rock dove and domestic breed clades are short, indicating low levels of differentiation among the populations, with the exception of the most basal branch where the West African rock doves are placed (Figure S4).

**Figure 3.**
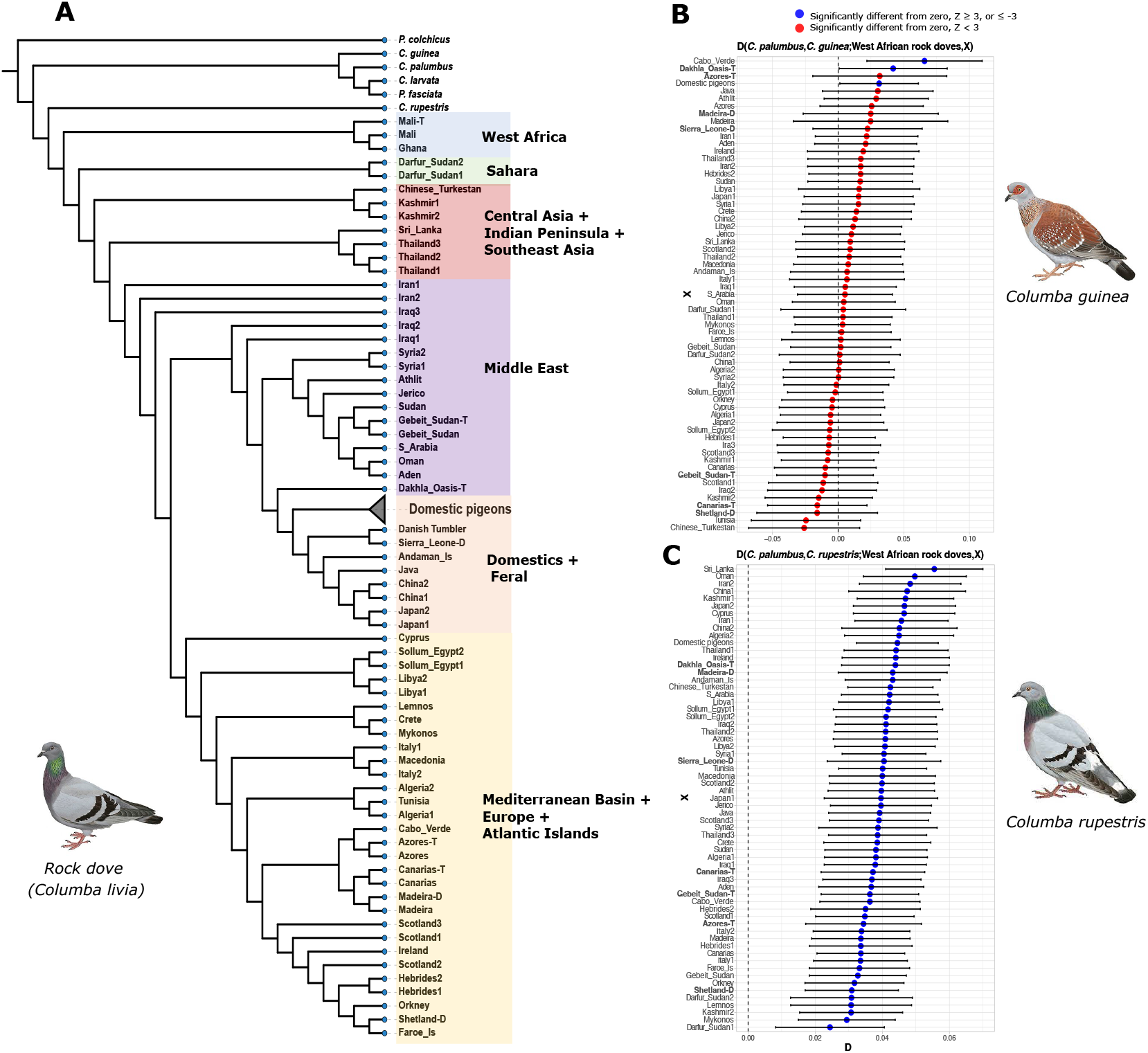
Phylogenetic relationships among wild and domesticated pigeons. A) Neighbor-Joining tree based on genomic pairwise-distances. Branch length was disregarded in the tree representation (see also Figure S4). Different colours indicate the main geographical regions. The domestic pigeon breeds are shown as a collapsed clade. B-C) *D-statistics* analyses testing the possibility of hybridization in the West African rock doves that could explain its basal placement in the tree. B) *D-statistics* test in the form *D*(*C. palumbus, C. guinea*; West African rock doves, X), where *C. palumbus* has been used as an outgroup, all West African rock doves are grouped in a single population, and X represent all other rock doves and domestic pigeons in the dataset. C) *D-statistics* of the form *D*(*C. palumbus, C. rupestris*; West African rock doves, X) testing for hybridisation between rock doves and its sister species *C. rupestris*. Horizontal bars show 3 standard errors estimated through a block jackknife approach. Tests with a resulting |*Z-*score|>3 were considered statistically significant (blue).

Rock doves are grouped into clades that clearly reflect their geographic origin (Figure 3A). The West African rock doves are placed as basal to all other rock doves, followed by the rock doves from the Sahara (Darfur, Sudan), Central Asia and Kashmir, Sri Lanka and Thailand, the Middle East and Red Sea region, and a clade including the Mediterranean Basin and Atlantic islands. In the estimated tree, all domestic breeds appear as monophyletic and closely related to the Middle East and Red Sea rock doves clade.

The historical feral pigeons form a clade together with the domestic pigeons, with the exception of the sample Madeira_D, that appears among the other Atlantic Islands rock doves. The modern feral pigeons from the USA are closely related to the domestic pigeons as it was reported in a previous study by Shapiro et al. 2013. One notable observation is that the *C. l. dakhlae* type specimen is placed basal to all domestic and feral pigeons. The Near East, Red Sea and Arabian peninsula rock doves are clustered together, and form the sister clade of the domestic and feral pigeons. Basal to these last mentioned sister clades, we find the samples Iraq1 and Iraq2 (Figure 3A). Also interesting is the placement of the rock doves from the Atlantic Islands, which are located within the Mediterranean Basin’s clade. Rock doves from the Atlantic Islands are divided in two smaller clades, North Atlantic islands rock doves, and Subtropical Atlantic islands rock doves, where the *C. l. atlantis* and *C. l. canariensis* types specimens are placed. Finally, the Atlantic Islands rock dove clade is placed as a sister clade of the rock doves from northwestern Africa, Algeria and Tunisia (Figure 3A).

Complementary to the NJ tree, we estimated a maximum-likelihood phylogeny summarising 1,000 independent trees using randomly chosen genomic regions of 5,000 bp each. Overall, the estimated tree topology is similar to the NJ tree and shows short internal branch lengths (Figure S5). The specimens from West Africa are again placed at the base of the rock doves, and this partition is supported by a high bootstrap value. Differences with the internal structure of the NJ tree could be produced by the distinct admixture levels among the rock dove populations, and domestic/feral pigeons. Despite these differences and the low bootstrap values, several relevant similarities are apparent, not least that all domestic pigeons appear monophyletic, and are closely related to the feral birds, with the specimen Syria1 being at the base of this clade. With regards to the differences between the NJ and maximum-likelihood approaches, we find that the placement of the Saharan rock doves from Darfur, Sudan do not appear as basal, and that previously observed geographic clades appear fragmented and mixed with rock doves from diverse geographical origins, indicating low divergence in the genomic regions used for build the phylogeny and possible admixture events with domestic lineages (Figure S5).

### Extensive introgression between C. rupestris and the rock dove

Our phylogenetic results show the rock doves from western Africa (Mali-T, Mali, and Ghana) are basal to all other rock doves in the dataset, which is unexpected given the proposed origin of the species in Asia (Johnston and Janiga 1995). To test the possibility of hybridisation between the rock doves from western Africa and other species from the *Columba* genus as an explanation for their unexpectedly divergent basal position in the tree, we used *D-statistics*. Of particular interest was the possible hybridisation with the speckled pigeon *C. guinea*, which both overlaps in distribution with the West African rock dove populations, and shares with them the phenotype of a highly developed red orbital ring (Baptista et al. 2009). We used *D-statistics* of the form *D*(*C. palumbus, C. guinea*; West African rock doves, X), where *C. palumbus* is used as outgroup, all the West African rock dove specimens are grouped into a single population, and X represents all rock dove, feral pigeons and domestic pigeons in our dataset. In all *D-statistics*, domestic pigeons were grouped as a single population according to the previous results, and the *qpWave* analysis that confirmed the close relationships among the pigeon breeds (Figure S6). If genomic elements of the West African rock doves derive from hybridisation with *C. guinea*, we would expect to obtain significant negative *D* values, indicating that the West African rock doves share more genetic drift with *C. guinea* than with the rest of the rock doves tested in the analysis. The results of the analysis however provide no evidence in support of this hypothesis. The only statistically significant values that indicate shared genetic drift with *C. guinea* involve the domestic pigeons, as well as the specimen from Cape Verde and the *C. l. dakhlae* type specimen from the Dakhla Oasis, both apparently closely related to domestic breeds according to our previous results (Figure 3B). Patterns of admixture between *C. guinea* and domestic breeds were reported in Vickrey et al. 2018. Continuing with the exploration for possible hybridisation events between species from *Columba* genus and rock doves, a second *D-statistics* analysis was performed using the hill pigeon *C. rupestris*, as a source population of the form *D*(*C. palumbus, C. rupestris*; West African rock doves, X). Interestingly, these results indicate a clear signal of shared genetic drift between *C. rupestris* and all samples in X, with the exception of the West African rock doves (Figure 3C).

### Domestic pigeons are closer to wild populations from the Middle East

The exact geographic location of the rock dove’s domestication reaims debated. Our phylogenetic results show domestic pigeons form a clade that is most closely to the Middle East clade (that includes samples from the Red Sea region), and the Mediterranean Basin-Europe-Atlantic Islands clade (Figure 3A). In order to assess this question, we implemented a set of *D-statistics* analyses of the form *D*(*C. palumbus*, Middle East rock doves; Mediterranean Basin-Europe-Atlantic Islands rock doves, Domestic pigeons) and the alternative that switches the H2 and H3 populations, *D*(*C. palumbus*, Mediterranean Basin-Europe-Atlantic Islands rock doves; Middle East rock doves, Domestic pigeons). In doing so we were able to test the tree structure and try to detect signals of gene flow with indications of directionality (Figure 4). In this analysis, *C. palumbus* was used as an outgroup, each historical rock dove from each clade was tested individually, and domestic pigeons were grouped in a single population. We expect that the *D-statistics* result would be statistically non-significant when the correct tree structure is tested and, if the incorrect structure is tested, the result will be positive and statistically significant, indicating high levels of shared genetic drift due a more recent common ancestry. The results show that when Middle East rock doves were used as population H2, the Z-score tends to be statistically non-significant, particularly when comparing specimens from Syria, Palestine (Althlit and Jerico) and the specimen Iraq1 with the rock doves from the Mediterranean Basin, Europe and the Atlantic Islands. In contrast, the values tend to become positive and statistically significant for the rest of the rock doves from the Middle East clade. When Middle East rock doves were used as population H3, the obtained Z-score values for the specimens Syria1, Syria2, Althlit, Jerico and Iraq1, are especially high, positive and statistically significant when compared with rock doves from Mediterranean Basin, Europe and the Atlantic Islands. In contrast, for the rest of the Middle East clade rock doves the tendency of the Z-score values is to decrease and become statistically non-significant. As can be observed in the Figure 4, the non-overlapping Z-score distribution patterns obtained when testing the rock doves Syria1, Syria2, Althlit, Jerico and Iraq1 suggest that these are the most closely related populations to the domestic breeds. An overlapping Z-score distribution is obtained when the rest of the Middle East rock doves are tested (Aden, Oman, Iraq2, S_Arabia, Gebeit_Sudan-T, Gebeit_Sudan and Sudan) as H2 and H3 populations, reflecting a less clear relationship between domestic pigeons, and a possibly higher degree of gene flow among all the different tested populations.

**Figure 4.**
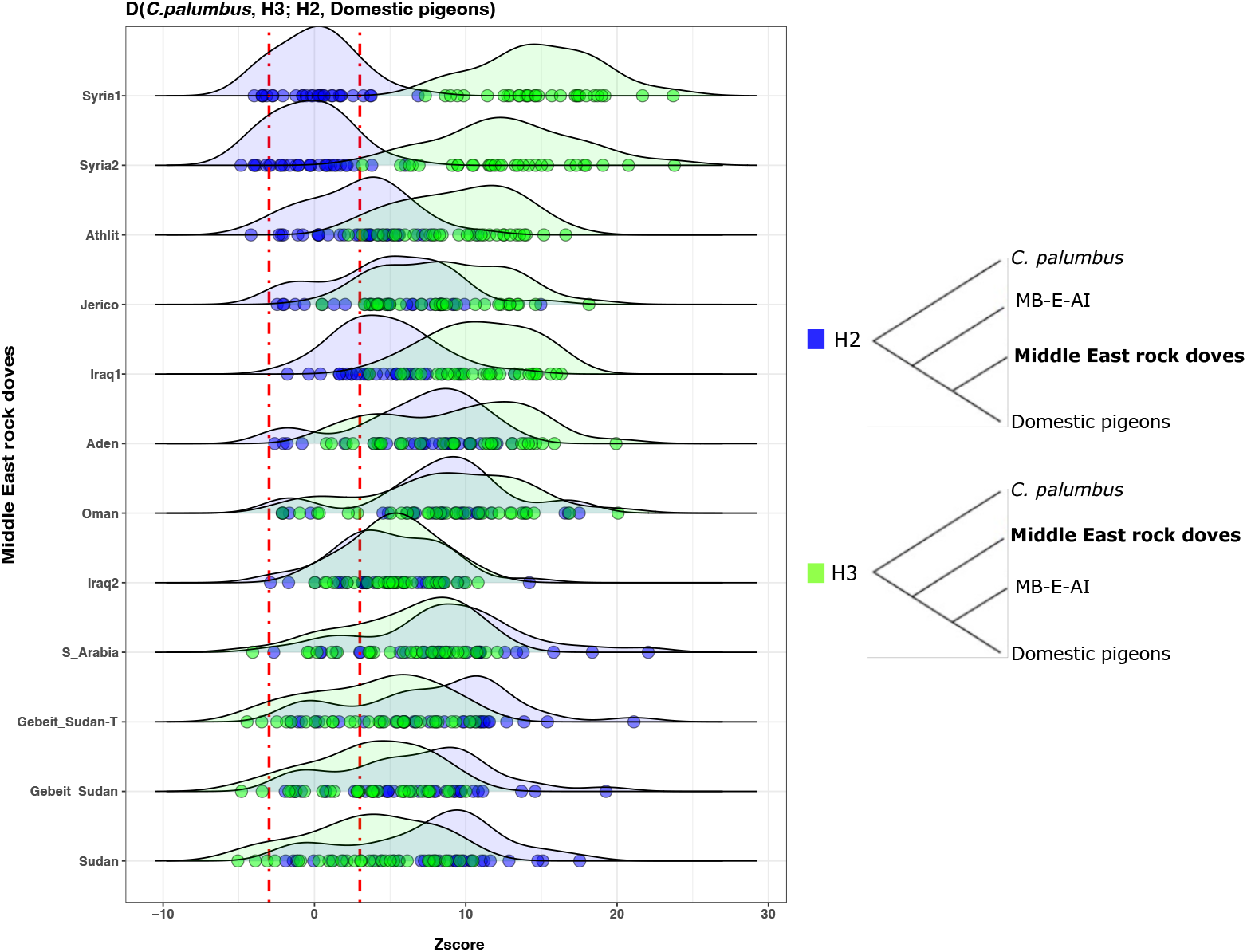
Evaluating the position of the domestic pigeon breeds in the tree using *D-statistics*. Density plots showing the Z-scores distributions obtained from the *D-statistic* analyses testing the tree topology between the: domestic pigeons, Middle East rock doves, and rock doves from the Mediterranean Basin-Europe-Atlantic Islands (MB-E-AI). The tests were performed for each sample individually, except for the domestic pigeons that were grouped in a single population. The two tree topologies tested are shown as diagrams in the right side of the figure. When Middle East rock doves were tested as an outgroup to the domestic pigeons and MB-E-AI rock doves, Z-scores were overall more positive and statistically significant, mainly for populations from Syria, Palestine and Iraq. In contrast, when Middle East rock doves were placed as forming a clade with the domestic pigeons, Z-scores values were lower and a higher proportion was non-statistically significant. Dots represent the Z-score obtained for each MB-E-AI specimen tested. Red dotted lines in the x axis show the range for non-statistically significant values (|Z|< 3).

### The type specimen C. l. dakhlae has an admixed origin

Our dataset allowed us to investigate the ancestry of the *C. l. dakhlae* type specimen (Dakhla_Oasis-T). The *C. l. dakhlae* subspecies is restricted to the Dakhlae and Kharga Oases in Central Egypt, and is characterized by its very light homogeneous colouration (Urban etl al. 2014; Dickinson and Remsen 2013; Cramp 1985). Our results show a close relationship of the *C. l. dakhlae* type specimen to the domestic breeds. We implemented a *D-statistics* analyis to decipher if this closeness is due to (i) hybridisation with domestic breeds, (ii) because it is ancestral to domestic breeds, or (iii) it is an ancient domestic lineage by itself. In this case, the *D-statistics* test was performed in the form *D*(*C. palumbus*, X; Dakhla_Oasis-T, Domestic pigeons), where *C. palumbus* was used as outgroup, X represents all historical rock doves in the dataset, and the domestic pigeons are grouped in a single population. If Dakhla_Oasis-T belongs to the ancestral lineage of all domestic breeds or to an ancient domesticated lineage, we would expect to obtain non-statistically significant *D* values. Alternatively, if the Dakhla_Oasis-T is a hybrid between domestic breeds and wild rock doves, the obtained *D* values would deviate from zero, and be statistically significant, showing different levels of affinity between the historical rock doves and the domestic pigeons. The results seem to point to that the Dakhla_Oasis-T is admixed, sharing more genetic drift with rock doves from similar geographic regions, mainly with specimens from the Red Sea region (S_Arabia, Aden, Sudan, Gebeit_Sudan, Gebeit_Sudan-T and Oman), when compared with the domestic pigeons. But, for the rest of the cases, the *D* values were non-statistically significant, suggesting a higher affinity between the Dakhla_Oasis-T and domestic pigeons than with the other rock doves in the dataset (Figure S7).

## Discussion

### Reconstructing the evolutionary history of rock doves through a hypothetical scenario based on the Pleistocene refugia theory

It has previously been proposed that the rock dove’s geographic origin is in Asia, based on the distribution of its closest sister species, *C. rupestris* (Johnston and Janiga 1995). This is striking in light of the fact that our results clearly indicate that rock doves from West Africa are both divergent and basal to all other rock doves included in this study. As our analyses enable us to discount the hypothesis that this basal placement is caused by hybridisation of West African rock doves with *C. guinea*, we propose an alternate hypothesis to explain the observation, related to the effect of Pleistocene climate change on the Sahara and the Sahel regions.

Global climate fluctuations during the Pleistocene glacial periods, produced drastic changes in the distribution and abundance of flora and fauna, leading to fragmented species distributions, population reductions, high extinction rates, but also, speciation events (Nadachowska-Brzyska et al. 2015; Hewitt 2000, 1996; Lovette 2005; Weir and Schluter 2004; Le Houérou 1997; Mayr and O’Hara 1986). As mentioned by Hewitt (2000), the current genetic structure of populations and species has been mainly formed during the Quaternary Ice Ages, which generated populations with allopatric distributions and subsequent genetic differentiation. In the case of the avifauna, several studies have described the changes in population size along the Pleistocene. In particular, the Last Glacial Maximum (LGM) was an important driver of speciation in boreal superspecies, and to a smaller degree, in superspecies from subboreal and neotropical regions of the Americas (Weir and Schluter 2004). Such impact in shaping population structure is likely to have affected other regions of the world.

During the Quaternary period, the Sahara and the Sahel underwent several dry and wet cycles associated to the glacial and interglacial periods (Manning and Timpson 2014; Larrasoaña et al. 2013; Le Houérou 1997), with an estimate of 8 to 10 arid-wet periods during the last 125,000 years alone (Le Houérou 1997). During the LGM, ca.18,000 years ago, the Sahara region had an hyperarid period, where the desert conditions extended south to the modern Sahara limits (Larrasoaña et al. 2013), and today West African forested areas were covered by open woodlands and grasslands with a semi-arid climate (Nichol 1999). This hyperarid period was preceded and followed for what is known as Green Sahara Periods (GSPs) (Larrasoaña et al. 2013).

In light of this context, we propose a hypothetical scenario that could explain the apparent contradiction of West African rock doves falling basal to all other rock doves, if the species originated in Asia: 1) Previous to the LGM, *C. livia* had a similar continental distribution to the modern wild members of species, but with a more widespread distribution in the Sahara region due to the favorable wet conditions at that time. 2) During the LGM, the southward advance of the ice sheet in the northern latitudes, and the hyperarid condition in the Sahara and the Sahel, drastically diminished the rock dove populations worldwide, reducing it to two main allopatric populations that survived in more stable or favorable environment for the species: one in West Africa, and a second in Central Asia. 3) During this same period of time, the Central Asian rock dove population hybridised with *C. rupestris*, as suggested by the *D*-statistic tests (Figure 3C). 4) After the LGM, the new favorable environmental conditions allowed the Central Asian rock dove population to spread and recolonise the pre-LGM distribution range, including the Sahara region that was under the Holocene GSP (ca. 6,000-10,000 years ago), producing a secondary contact with the West African rock dove population, and a hybrid zone. 5) Finally, the current arid Saharan period led their habitat to either partially or totally split the West African rock dove populations from the rest of the African rock doves, yielding today’s distribution of the species.

The rock dove range distribution previous to the LGM is unknown, but taking into consideration the high dispersal capacity, and adaptability of rock doves, it would not be surprising if the species’ distribution was as wide as it is today. Certainly, the divergence and basal position of West African rock doves in our results suggests that the distribution reached the Western part of Africa. On the other hand, there is evidence of a decrease in the effective population size of rock doves during the LGM (*cf*. Nadachowska-Brzyska et al. 2015). It is possible that several populations survived in different Pleistocene refugia, as in the south of Europe, but our data does not show evidence of populations that diverged in allopatry other than the West African populations. This could mean that most populations became extinct due to its small size, inbreeding depression and accumulation of deleterious mutations (Nadachowska-Brzyska et al. 2015; Frankham 2005), or that they were diluted and replaced by the main rock dove post-LGM recolonization wave.

The *D-statistics* result showing that all rock doves used in this study (including domestic breeds) share more alleles than expected with *C. rupestris* when compared with the West African populations, suggests that this shared genetic signature among rock doves from very different locations was acquired by a common ancestry. Most likely, this hybridisation occurred in the ancestral rock dove population around the overlapping distribution between the two species in Central Asia (hybridisation between *C. rupestris* and *C. livia* has been reported by Johnston and Janiga 1995). In our hypothetical scenario, this same population spread after the LGM and gave rise to the modern rock dove populations in Asia, Europe, North Africa, the Sahara, East Africa, as well as the Atlantic Islands. The latter is supported by the low genetic diversity observed in the branch lengths of the estimated trees, as well in the MDS plot among the historical rock doves. It has been proposed that in the context of interglacial periods, rapid expansions in northern populations would produce increased homozygosity or low genomic diversity due to founder effects and continuous bottlenecks. Otherwise, southern populations in more stable habitats tend to diverge by repeated allopatry during several glacial cycles and by being protected by hybrid zones (Hewitt 2000; 1996), as could be the case of West African rock doves.

Pleistocene refugia theory typically predicts the establishment of a secondary contact after the expansion of the allopatric populations, and the development of hybrid zones (Mayr and O’Hara 1986). Our results indicate that rock dove specimens from Darfur, Sudan, show some evidence of such secondary contact and subsequent hybridisation. It is possible that the West African rock doves expanded to the east through the Sahara, and the Asian populations expanded toward African and then the Sahara, creating a secondary contact around Central Sahara that allowed the development of an hybrid zone. Once the Sahara became arid again, the hybrid zone disappeared and the populations became allopatric. It will be important to explore other rock dove populations from the Sahara to confirm the previous existence of an hybrid zone, and the complete or partial allopatry condition of the West African rock dove populations today.

Although our hypothesis aims to explain in a broad sense the rock dove’s evolutionary history, it will be necessary to explore in more detail and in a regional scale the rock doves’ diversity to test this hypothesis, and provide evidence that could contribute to the support or deny of each of the steps proposed here, as well as to add complexity to the description of the evolutionary history of rock doves.

### Insights into the domestication of the rock dove

Most authors have argued in support of the hypothesis that rock doves were independently domesticated at multiple locations in the Mediterranean Basin and the Near East (Shapiro and Domyan 2013; Johnston and Janiga 1995; Johnston 1992). Although today domestic breeds have low levels of genetic differentiation, this is likely due to recent intense crossing among breeds. As a consequence, resolving the timing and number of domestication events may be very challenging (Shapiro and Domyan 2013). Under a scenario of multiple domestication events, we could expect that different breeds would display different affinities with rock doves from distinct geographical locations, despite the high levels of crossing among breeds. In our results, domestic pigeons appear to be closely related and form a monophyletic clade in the NJ and maximum likelihood trees, the latter having considerably high support (bootstrap of 74%) in comparison to other internal branches in the tree. Even if high levels of crossing among breeds has happened, it is somewhat unexpected to see such a level of genomic homogenisation in all the breeds, effectively blurring other signals of ancestry across the different genomic regions. Interestingly, we do find that domestic pigeons are the sister clade of the Middle East rock dove populations, and the results of our *D-statistics* analyses indicate they are specifically close to populations from the Levante (Syria, Iraq and Palestine), which overlaps with the Fertile Crescent, the region that is believed to have been the center of origin of several other domesticated species, and has also been proposed to be the geographic origin of domestic pigeons (Driscoll et al. 2009).

Given this, our results point to at least one domestication event in the Levant region that gave rise to all domestic breeds analysed in this study. Although we cannot rule out the possibility of several domestication events, our data provides no evidence that domestication involved multiple genetically very distinct populations. We believe that rock dove synanthropy was common across a big part of its natural distribution, but complete domestications might be less common than previously assumed.

### Taxonomic implications

The intraspecific taxonomic classification of rock doves has been established on clinally distributed phenotypic characters that intergrades among populations such as coloration, plumage patterns, as well as body size (Murton and Clarke 1968). The latter, leads to an oversimplification of their diversity and complicates defining the distribution range of each subspecies (Johnston 1992). Since the 19th and early 20th centuries, when the rock dove subspecies were defined, few major taxonomic changes have been done, mainly, invalidating some subspecies due to their similitude with other adjacent subspecies (Gibbs et al. 2001; Cramp 1985; Murton and Clarke 1968).

The rock doves from West Africa are distributed in Mauritania, Mali, Ghana, Senegambia and Guinea (Urban et al. 2014), and are currently recognised as *C. l. gymnocyclus* Gray, 1856. This subspecies is characterised by its smaller size and very dark color in comparison to the nominate *C. l. livia* Gmelin, 1789, and the extended bright scarlet orbital skin (Gibbs et al. 2001). Although Mali populations were initially separated based on their smaller size and paler plumage into the subspecies *C. l. lividor* Bates, 1932, they have subsequently been considered a synonym of the *gymnocyclus* subspecies (Gibbs et al. 2001). According to the close relationship shown in our results between the specimens from Mali, including the type specimen for *C. l. lividor* (Mali-T), and specimens from Ghana, we believe that including *C. l. lividor* within *C. l. gymnocyclus* is correct.

Subtropical Atlantic Islands rock doves have been historically classified into two subspecies, the populations from the Canary Islands as *C. l. canariensis* Bannerman, 1914, and the populations from Azores, Cape Verde and Madeira as *C. l. atlantis* Bannerman, 1931. The *C. l. canariensis* subspecies has been described as smaller and darker than the nominate, but because the differences are considered very small, those populations are usually included in *C. l. livia* (Cramp 1985). On the other hand, *C. l. atlantis* is thought to have derived from feral pigeons due its high variability in plumage patterns and coloration (Gibbs et al. 2001; Cramp 1985; Muton and Clarke 1968), or from melanistic mutants from the Canary Islands (Cramp 1985). According to the observed genomic affinities in our results, rock doves from the Subtropical Atlantic Islands (including types specimens of *C. l. atlantis*, and *C. l. canariensis*) are closely related to populations from northwest Africa (Algeria and Tunisia) and the North Atlantic Islands, indicating a natural origin for these Subtropical Atlantic Islands rock doves. However, signals of admixture can be observed mainly in the specimens from Cape Verde and Azores (Figure 2C), which could explain the morphological variation that characterise these populations. Given the stronger affinities of the subtropical Atlantic Islands rock doves with other *C. l. livia* populations, we believe that, despite the signals of genetic introgression from feral birds, the original populations from the subtropical Atlantic Islands could belong to *C. l. livia* subspecies. The description of *C. l. atlantis*, and *C. l. canariensis* subspecies was probably based on specimens belonging to populations with high levels of admixture with feral pigeons.

The northeast Sudan rock dove populations that were described as *C. l. butleri* Meinertzhagen, 1921, are commonly included in *C. l. targia* Geyr von Schweppenburg, 1916 (Gibbs et al. 2001), in *C. l. gaddi* Zarudny and Loudon, 1906 (White 1965), or in *C. l. schimperi* Bonaparte, 1854 (Cramp 1985). According to the different analyses performed that included the type specimen of *C. l. butleri*, the rock doves from the Sudan Red Sea Province are more closely related to populations from the Arabian peninsula and, in a second level, to rock doves from the Levant region that belong to the subspecies *C. l. palestinae* Zedlitz, 1912. Even though we lack *C. l. schimperi* specimens from Egypt, our results suggest that *C. l. butleri* is closer to *C. l. schimperi* than to *C. l. targia* or *C. l. gaddi*. Vauri (1965) previously suggested the inclusion of *C. l. butleri* in *C. l. schimperi*. Therefore, based on our genomic data, we believe *C. l. butleri* is a synonym of *C. l. shimperi*.

An interesting case is the rock dove subspecies *C. l. dakhlae* Meinertzhagen, 1928 that is restricted to the Dakhla and Kharga Oases in Central Egypt, and is characterised by a very light pale plumage. Throughout our analyses, the Dakhla_Oasis-T specimen showed a close affinity to the domestic breeds. The *D-statistic* results indicated that this specimen has genetic components from domestic breeds and rock doves from nearby regions, mainly from the Red Sea, Arabic Peninsula, the Levante, North Egypt, and Sudan. It seems likely therefore that *C. l. dakhlae* derived from the hybridisation between feral pigeons and wild populations. Consequently, we argue that the *C. l. dakhlae* subspecies status is invalid due to its apparent feral origin. Nevertheless, we believe that rock doves from Dakhla and Kharga Oases offer an important opportunity to study the long term effects of domestic gene introgression in wild populations.

Of more relevance is that our genomic-based analyses suggest that rock doves of the *C. l. gymnocyclus* subspecies form the basal sister clade of all other rock doves, due to a long history of allopatry which derived in the divergence of its populations. For that reason, we believe that a taxonomic arrangement can be necessary to reflect their evolutionary history. We propose that *C. l. gymnocyclus* Gray, 1856 should be considered a full species, *Columba gymnocyclus* Gray, 1856, with *C. l. lividor* Bates, 1932 being a junior synonym.

Finally, the geographically distributed diversity among the historic rock doves (excluding West African populations) in our study, as well as the low differentiation among populations, suggest that rock doves form a continuum throughout their distribution range, which match with the clinically distributed phenotypic characters that have been used to previously criticise the rock dove subspecies classification. In the Johnston’s extensive (1992) study based on the measurement of several morphological characters of 222 male rock dove specimens from most of the historical distribution of the species, he concluded that some of the main subspecies (*C. l. gaddi, C. l. palestinae, C. l. livia, C. schimperi*, and *C. l. neglecta*) were arbitrarily distinguished in the borders of their distributions, concluding that those subspecies “have little potential evolutionary novelty in the absence of geographic isolation”. The genomic data indicates that the rock dove subspecies are difficult to support and identify from adjacent populations. Although we are missing specimens from intermedial geographical locations among some populations, we think their inclusion would strengthen the observed patterns. From our point of view, if *C. l. gymnocyclus* is raised to species status, *C. livia* can be considered as monotypic, or alternately, two subspecies can be recognized considering the rump colouration that has been used to differentiate rock dove populations: the nominate *C. l. livia* Gmelin, 1789, which presents a white rump colouration (Johnston and Janiga 1995), and the rest of the populations (which present mostly grey and occasionally white rumps) (Johnston and Janiga 1995) together into *C. l. intermedia*, Strickland, 1844, respecting the oldest subspecies name.

## Materials and Methods

### Dataset

We obtained a total of 64 dried toe pads and one alcohol-preserved toepad from historical *Columba livia* specimens dated from 1865 to 1986 and housed at the Natural History Museum at Tring, UK (Table S1). The specimens represent all currently recognised subspecies. The collection locations of these specimens cover most of the reported historical distribution of the species, including the Atlantic Islands, the Mediterranean Basin, the Saharan and West sub-Saharan Africa, Middle East, Central Asia, Sri Lanka, and Thailand. Eight of them are described as feral pigeons from China, Japan, Sierra Leone, Madeira Island, Java, and Andaman Islands. Among these specimens 3 are from Darwin’s collection, and 5 are type specimens of different taxa: *C. l. atlantis* Bannerman, 1931, *C. l. canariensis* Bannerman, 1914, *C. l. butleri* Meinertzhagen, 1921, *C. l. dakhlae* Meinertzhagen, 1928, and *C. l. lividor* Bates, 1932. The rest of the samples, according to their taxonomic description, belong to the currently accepted subspecies (Table S1).

To complement our historical dataset, we included 41 reference pigeon genomes (Shapiro et al. 2013), 39 of them from different domestic pigeon breeds, and 2 feral pigeons from the USA. Additionally, 6 outgroups were added to the final dataset: 1 common pheasant (*Phasianus colchicus*) (Liu et al. 2019), 1 band-tailed pigeon (*Patagioneas fasciata*) (Murray et al. 2017), 1 cinnamon dove (*Columba larvata*), 1 common wood pigeon (*Columba palumbus*), 1 speckled pigeon (*Columba guinea*) (Vickrey et al. 2018), and 1 hill pigeon (*Columba rupestris*) (Shapiro et al. 2013) (Table S2).

### Laboratory procedures

The historical samples were processed under strict clean laboratory conditions at the Globe Institute, University of Copenhagen. DNA extractions were performed using a digestion buffer for keratin following Campos and Gilbert (2012).The samples mRD_1 - mRD_10 were digested in 1ml buffer and purified using a binding reservoir as in Dabney et al. 2013, combined with Monarch columns and a binding buffer as in Allentoft et al. 2015.

Samples mRD_11 - mRD_65, followed the same overall approach. Tissue for each sample was was digested overnight in 0.3ml of the aforementioned digestion buffer., which was purified directly over Monarch DNA Cleanup Columns (5 μg) (New England Biolabs) using a 10:1 ratio of binding buffer (a modified version of Qiagen’s PB buffer) to digestive mixture, repeated 3 times. PE buffer (Qiagen) was then used to wash the sample, followed by a dry spin to remove any remaining residue, and finalised with two consecutive elutions to increase DNA yield, each of 15 μL of EB (Qiagen) buffer incubated at 37 C for 10 minutes.

Double-stranded DNA sequencing libraries were constructed following BEST 2.0 (Carøe et al. 2018) although with modifications to allow sequencing using BGI sequencing technology (Mak et al. 2017). Libraries were prepared using 32 μL of DNA in a final reaction volume of 80 μL. A maximum of 10μL of each double stranded DNA library template was used for one round of PCR amplification of 25 thermal cycles of: 30s at 95ºC (denaturation), 30s at 60ºC (annealing) and 110s at 72ºC (extension). Each 50 μL reaction contained 2.5U PFU Turbo CX

Polymerase, 1x PFU Turbo buffer, 0.4 mg ml-1 bovine serum albumin (BSA), 0.25 μM mixed dNTPs, 0.1 μM BGI forward index-primer, 0.1μM BGI reverse index-primer. Amplified PCR products were purified using a 1.4X ratio of HiPrep PCR clean-up beads (Magbio Genomics), were eluted in 28μL of Elution Buffer with TWEEN (0.05%). Purified, amplified libraries were then quantified using the Qubit 2.0 Fluorometer (ThermoFisher Scientific, Inc) and Fragment Analyzer (Agilent Technologies) and were sequenced across eight 100bp PE DNBseq-G400 lanes (BGI Europe).

### Data processing

PALEOMIX v.1.2.13.4 BAM pipeline (Schubert et al. 2014) was used to process the generated sequence reads and map them against the *Columba livia* reference genome (Cliv_2.1) (Holt et al. 2018). In brief, this pipeline includes the following steps: 1) trimming of sequencing adapters, using AdapterRemoval v.2.2.0 (Schubert et al. 2016) discarding reads shorter than 25bp after trimming (*--minlength* default value (25bp)); 2) sequence alignment to the reference genome using BWA v.0.7.17 backtrack algorithm (Li and Durbin 2009), disabling the use of the seed parameter unmapped reads were discarded; 3) identification and removal of PCR duplicates using Picard’s MarkDuplicates (http://broadinstitute.github.oi/picard/); 4) local realignment around indels using GATK v.3.8.3 IndelRealigner module (McKenna et al. 2010). The three most distant outgroups in our dataset (*P. colchicus, C. palumbus, C. guinea*) were mapped using BWA *mem* algorithm instead of *backtrack* given the reference data consisted of longer reads and we expect a higher proportion of mismatches to the reference genome due to their longer evolutionary distance..

### Characterization of the DNA damage patterns in the historical genomes

We used mapDamage v.2.0.9 (Ginolhac et al. 2011) with default parameters to estimate the patterns of nucleotide misincorporations and DNA fragmentation in the historical specimens and characterise their level of DNA damage (Table S1). In most cases we do not observe a substantially high proportion of cytosine deamination at the end of the reads. This is consistent with previous observations showing historical specimens do not always show high C to T damage patterns (Sánchez-Barreiro et al. 2021). However, in order to avoid any potential bias caused by the small proportion of damage in the sequencing data, we restricted the analyses to transversion sites.

### SNP calling

Due to the heterogeneity in the average depth of coverage obtained from our historical samples, we used a pseudo-haploid calling approach instead of performing genotype calling. We used ANGSD v.0.931 *-dohaplocall* 1, which randomly samples a single base per site for every sample (Korneliussen et al. 2014). We selected sites with the following quality filters and parameters: *-doCounts* 1, *-minMinor* 2, *-maxMis* 10, *-C* 50, *-baq* 1, *-minMapQ* 30, *-minQ* 20, *- uniqueOnly* 1, *-remove_bads* 1, *-only_proper_pairs* 1, *-skipTriallelic* 1, *-doMajorMinor* 1, and *-GL* 2. Transitions were removed to avoid possible DNA damages in the historical rock doves and pigeon sequences. The SNPs calling was restricted to the first 128 scaffolds that are 1Mb or longer. We then used Plink v1.90 (Chang et al. 2015) to discard sites with a minor allele frequency below 0.01. The final SNP panel contained 1,642,881 transversion sites and 112 samples.

### Multidimensional scaling analysis

To explore the population structure in our dataset, a multidimensional scaling (MDS) analysis was implemented, allowing us to visualise the genetic structure in our samples. Pairwise-distances were estimated for the dataset described in the previous section using Plink v1.90. We estimated two MDS analysis, one including all samples (with exception of the outgroups) (Figure 2A), and a second excluding the divergent rock doves from the West Africa in order to observe in more detail the population structure among the rest of the genomes (Figure 2B and S1). All the results were visualised using the ggplot2 library in R (Wickham 2016).

### Admixture analysis

ADMIXTURE v.1.3.0 (Alexander et al. 2009) was used to estimate the ancestry components in our dataset, excluding the outgroup species. The pseudo-haploid dataset described in the previous section was used as input for this analysis. ADMIXTURE was run assuming from 2 to 10 (K={2…10}) ancestry components and 20 independent replicas for each K value were performed. The replicas with the best likelihood value were selected. We performed cross-validation for each value of K (Figure S2). R library Pophelper v.2.3.1 (Francis 2017) was used to visualise the admixture results.

### Neighbor-joining tree and maximum likelihood phylogeny

We estimated a neighbor-joining (NJ) tree based on a pairwise-distance matrix estimated from our pseudo-haploid dataset using Plink v1.90. The NJ tree was estimated using the R library ape (Paradis and Schliep 2019) and the Interactive Tree Of Life (iTOL) v4 online tool (Letunic and Bork 2019) was used to visualise it.

To infer a maximum likelihood phylogeny, we generated genomic consensus sequences for each genome using ANGSD v.0.931 (-*dofasta 2* option and applying quality filters *-minQ* 20 and *-minmapq* 20) and, using the *Columba livia* reference genome (Cliv_2.1). Then, 1000 independent phylogenetic trees were estimated in RAxML-ng v.0.9.0 (Kozlov et al. 2019) under the GTR+G evolutionary model, using 1000 random regions of 5000 bp taken from the previously created genomic consensus sequences. In the next step, all the gene trees were summarised to generate a species tree using ASTRAL-III (Zhang et al. 2018). The final tree was visualised using the Interactive Tree Of Life (iTOL) v4 online tool (Letunic and Bork 2019).

### qpWave

Taking in consideration the highly probable admixture between domestic pigeons and rock doves in our dataset, we decided to implement pairwise qpWave tests in ADMIXTOOLS v.5.1 (Patterson et al. 2012) to try to detect the minimum number of ancestral sources needed to explain the variation found in the domestic pigeon cluster. The test was done for all pairs of domestic pigeons (left populations) using the option ‘allsnps = YES’. The Right populations used in the test included *Columba palumbus* as the outgroup and samples representing the main clusters found in the NJ-tree: Mali, Darfur_Sudan2, Kashmir1, Thailand1, Iran1, and Iraq1. The analysis showed that most of the domestic pigeons can be grouped together, indicating their close relationships as was observed in the other obtained results (Figure S6).

### D-statistics

To evaluate the possible high levels of admixture between domestic and wild birds, we created a subset of our dataset including all historic samples and some domestic pigeons that represent the main genetic clusters identified in the qpWave analysis (Fantail, Indian fantail, Starling, Scandaroon, Lahore, English trumpeter, Racing, Oriental, Archangel, Cumulet) to perform SNP calling and try to avoid bias toward the domestic pigeons. For this purpose genotype likelihoods were estimated (-GL 2 -doMaf 2) using ANGSD v.0.931 restricting to SNPs with a p-value threshold of 1e-6. The selected sites were extracted from the first pseudo-haploid dataset described above, and a minor allele frequency filter of 0.01 was applied using plink.

This new pseudo-haploid dataset was used to explore admixture patterns and test the NJ-tree and phylogenetic tree topologies using *D-statistics* as implemented in ADMIXTOOLS v.5.1. When a test in the form D(Outgroup, A; B, C) deviates from 0, it suggests possible gene flow between A and B or A and C; if D < 0, A and B are sharing a higher level of genetic drift that expected, indicating possible gene flow; if D > 0, it indicates possible gene flow between A and C. Deviation from 0 was considered statistically significant when Z-score was under -3 or above 3. The significance of the test was estimated using a weighted block jackknife procedure over 1 Mb blocks. We performed the following tests:

1. To verify the basal position of the West African rock doves from Mali and Ghana, we tested the possibility of hybridisation with close related species *C. guinea* and *C. rupestris* in the form *D*(*C. palumbus, C. guinea*/*C. rupestris*; West Africa, X) where X represent all the other genomes in our dataset, including the domestic pigeons that were grouped in a single population, and the West African rock doves were also grouped in a single populations.
2. We implemented two complementary *D-statistics* tests to explore these observed relationships between domestic breeds and historical rock doves from the Middle East group. First, we performed a test of form *D*(Outgroup, MB-E-AI; Middle East rock doves, Domestic pigeons), where all domestic pigeons were grouped as one population, each sample from the Middle East region were tested and MB-E-AI represent each specimen from the Mediterranean Basin-Europe-Atlantic islands. Then we estimated a second test of the form *D*(Outgroup, Middle East rock doves; MB-E-AI, Domestic pigeons), switching the wild populations with the goal to confirm the relationships and obtain some clues for the gene flow direction.
3. A *D-statistics* test was performed to assess the close relationship of the *C. l. dakhlae* type specimen with the domestic breeds found in the different results. The form of the test was *D*(C.palumbus, X; Dakhale-Oasis-T, Domestic pigeons) where X represents all the historical rock doves in our dataset.

## Acknowledgments

Funding for this research was provided by ERC award Extinction Genomics and DNRF143 Center for Evolutionary Hologenomics. G.H-A. is supported by the Consejo Nacional de Ciencia y Tecnología from Mexico (CONACyT 576743).

## Data availability

Generated raw sequence reads have been deposited at the European Nucleotide Archive (Project ID: PRJEB61000).

## Author contributions

G.P. and M.T.P.G. conceived the study. H.v.G provided the samples. Funding was obtained by M.T.P.G. E.C., M.M.C., G.P., and M-H.S.S processed the samples in the laboratory. G.H-A. performed the data analysis with support from J.R-M. G.H-A wrote the manuscript with support from J.R-M., and S.G., H.v.G, and M.T.P.G. All authors revised the final manuscript.

