## Supplementary material for "Redefining rock doves, *Columba livia*, using historical whole genome sequences": Suplemental information


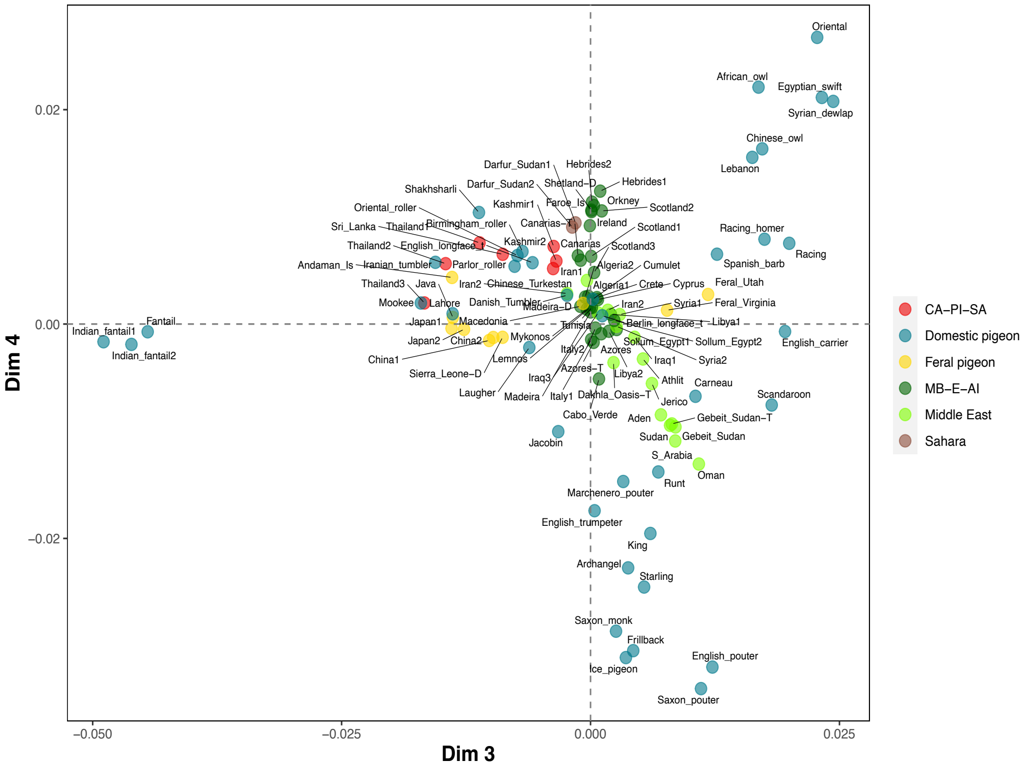


**Figure S1. Multimensional scaling plot. Related to Figure 2B.**
Multidimensional scaling plots (MDS) for dimensions 3 and 4 generated from pseudo-haploid data containing 1,642,881 transversion sites. West African rock doves were excluded. Colours represent the feral and domestic pigeons, as well as the geographic origin of the rock doves. Red colour includes samples from Central Asia, Peninsular India, and Southeast Asia (CA-PI-SA); dark green includes samples from the Mediterranean Basin, Europe, and the Atlantic Islands (MD-E-AI).

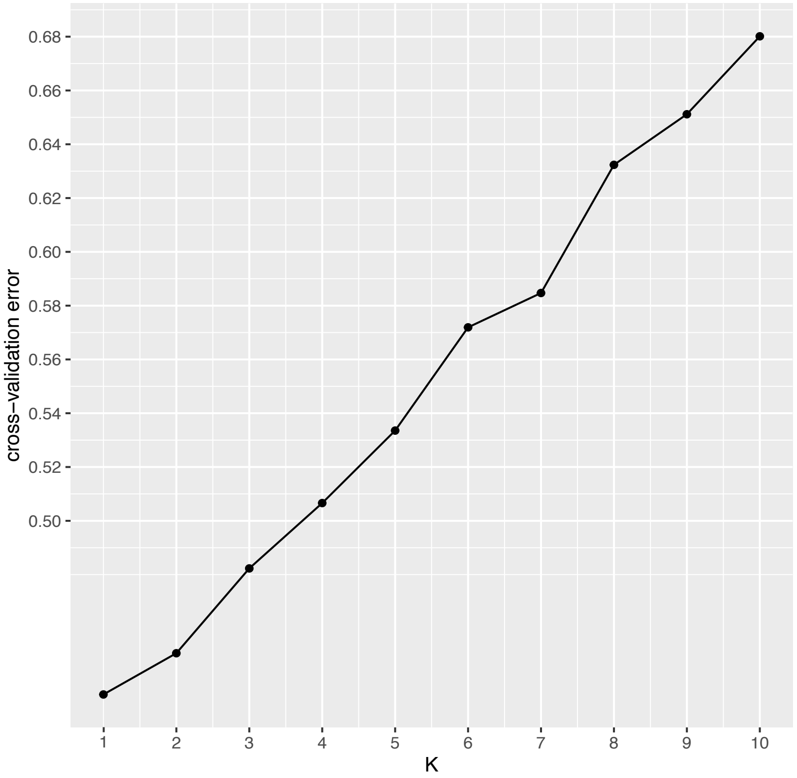


**Figure S2. Cross-validation errors plot.**
Cross-validation errors estimated in ADMIXTURE for 1 to 10 ancestral components (K). All rock doves in the data set were analyzed (historical rock pigeons, feral pigeons, and domestic pigeons), excluding the outgroups. Lower values of cross-validation errors are considered a better model fit. The lower obtained value corresponds to one ancestral component.

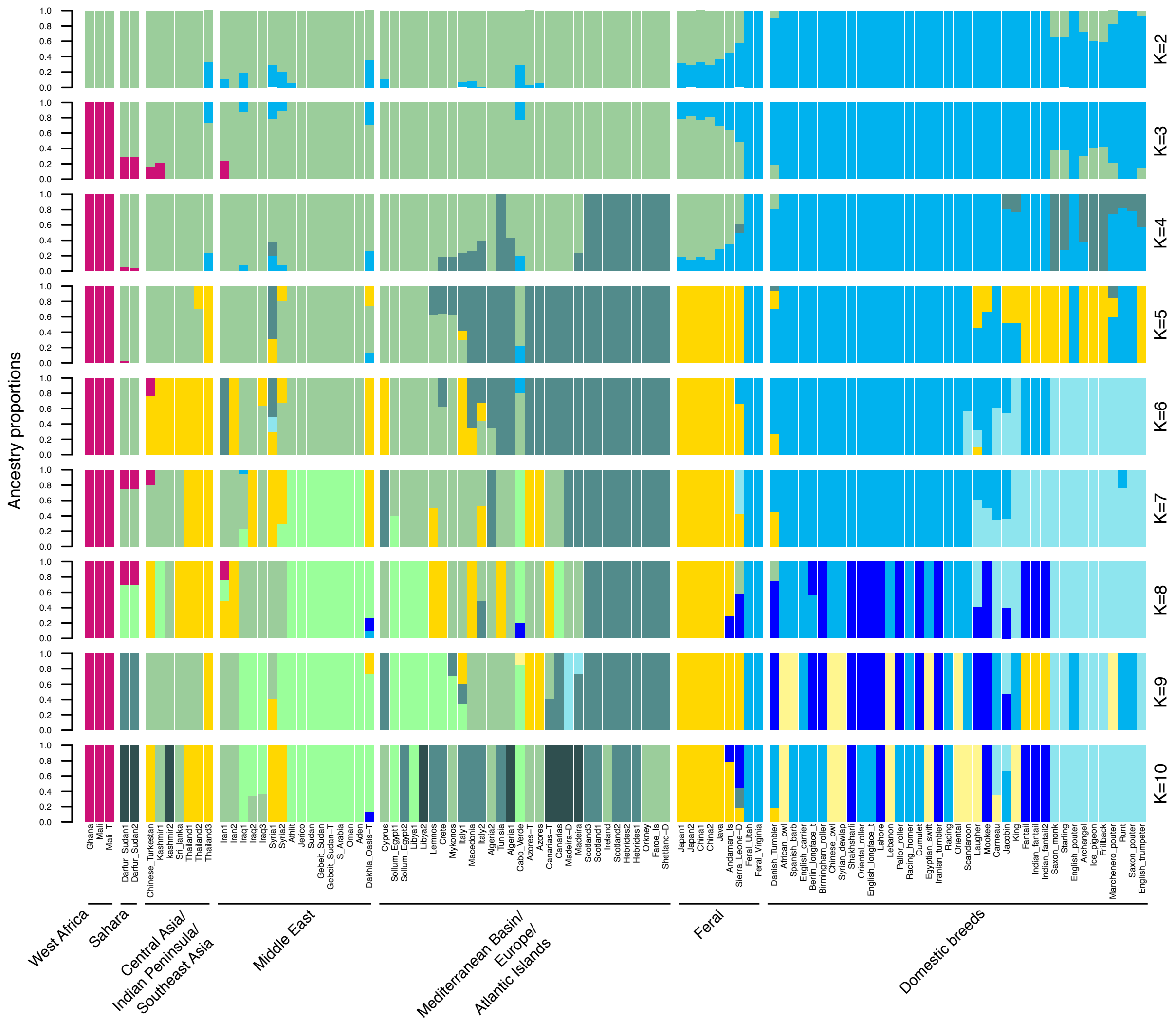


**Figure S3. Extended admixture graphs estimated for 2 to 12 ancestry components (K) in ADMIXTURE. Related to Figure 2C.**

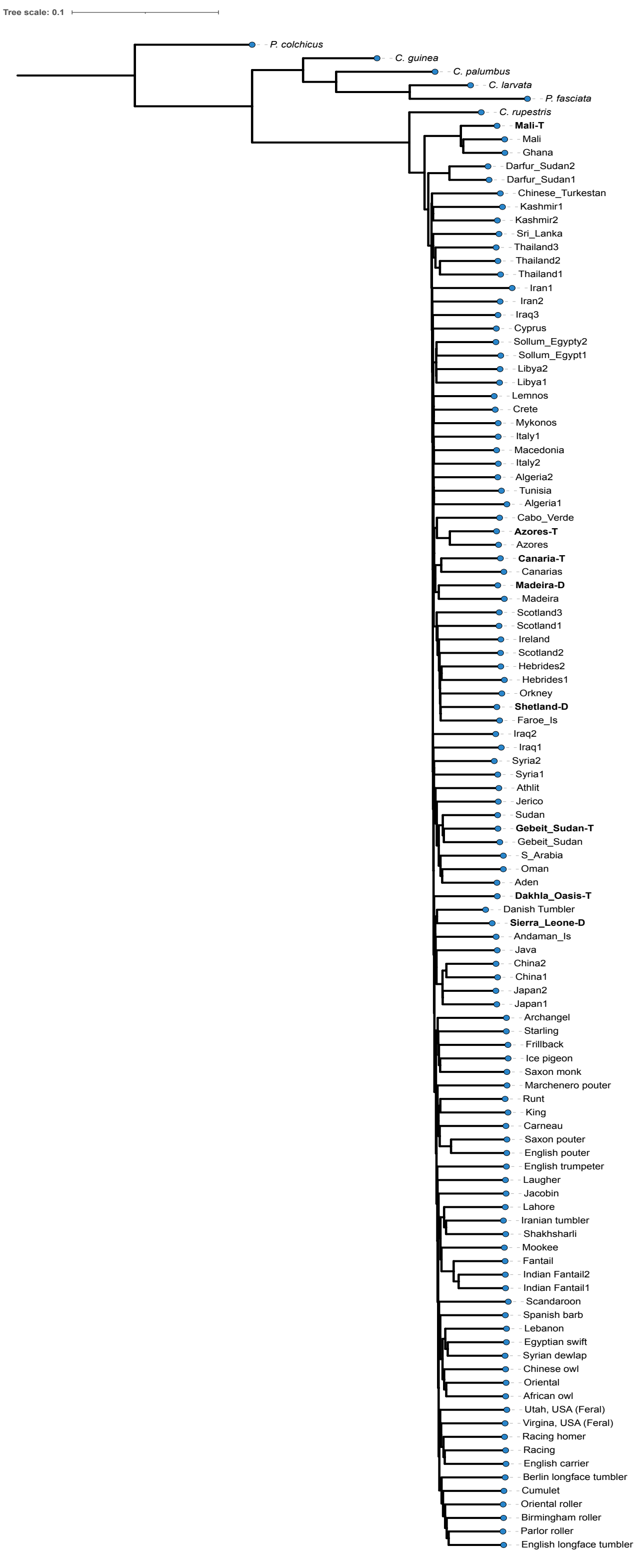


**Figure S4. Distance based Neighbor-Joining tree with branch lengths. Related to Figure 3A.**

Rock doves from Darwin's collection (D) and holotype specimens (T) are shown in bold.

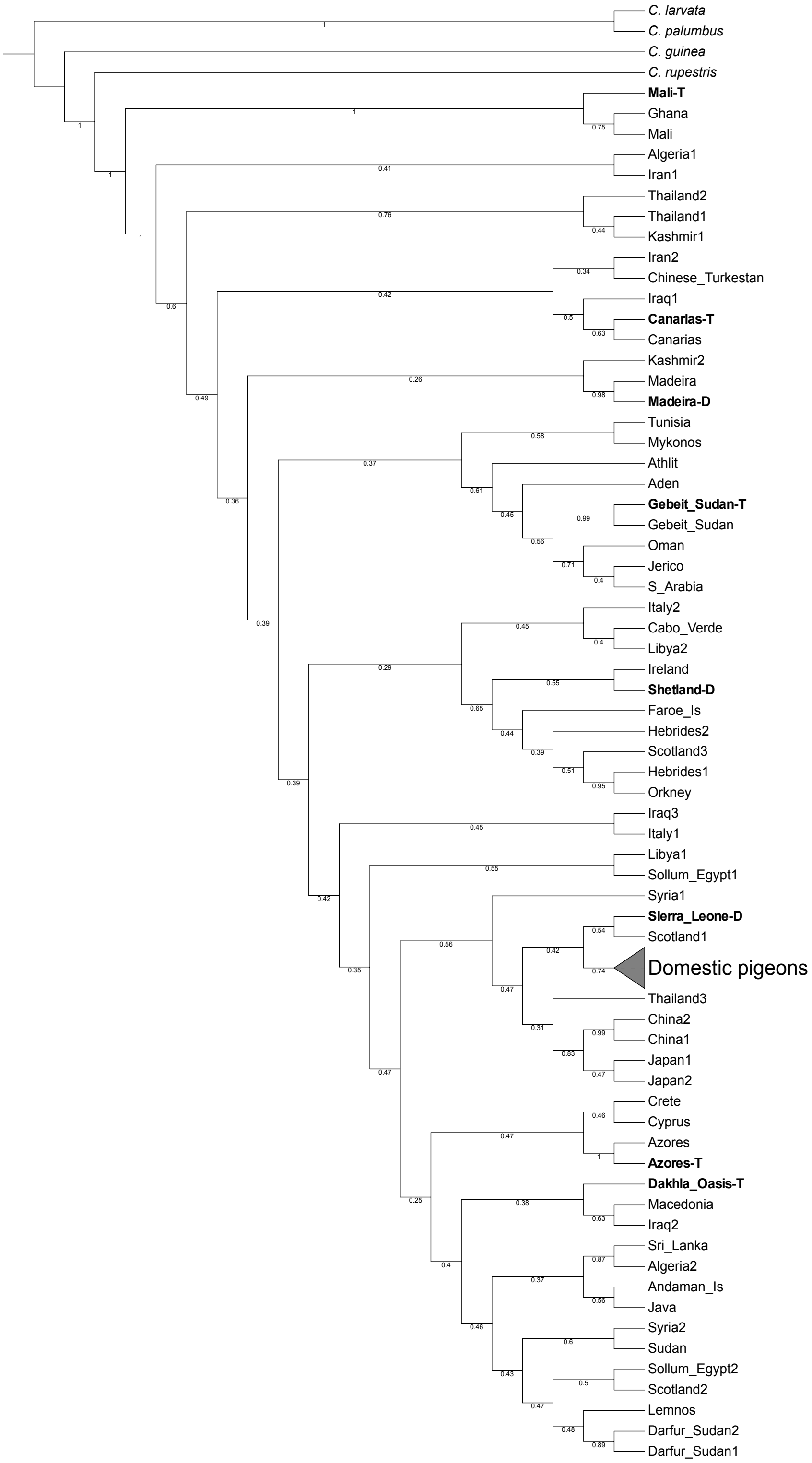


**Figure S5. Phylogenetic relationships estimated by genomic data.**
Maximum likelihood phylogeny built on 1000 concatenated trees. Bootstrap values are shown for each internal branch. Branch lengths were not used for a better visualization of the internal tree structure. Rock doves from Darwin's collection (D) and holotype specimens (T) are shown in bold.

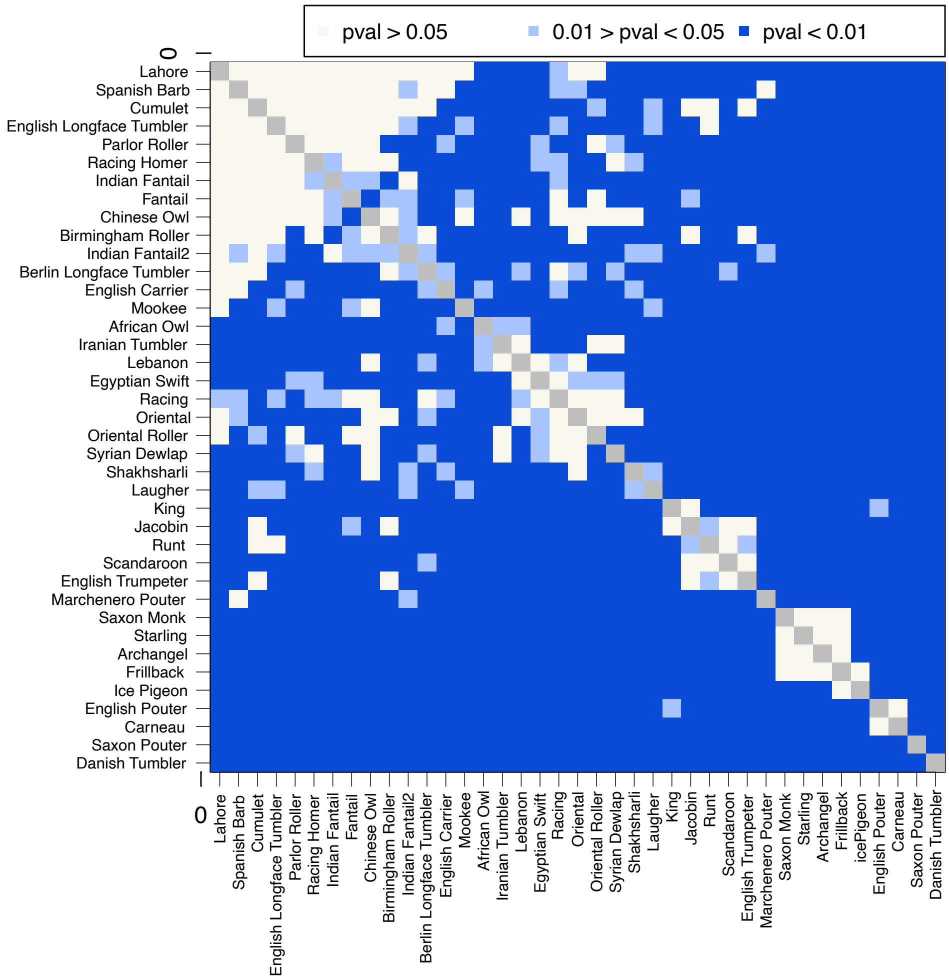


**Figure S6. Heatmap of pairwise qpWave p-values estimated for the domestic pigeons in the dataset.**
The plot shows the pairs of domestic pigeon genomes that can be modeled with the same ancestral component and can be identify as independent gene pools. The populations used to compare the pairs of domestic pigeons (Right populations) are *C. palumbus* as the outgroup and samples representing the main clusters found in the NJ-tree: Mali, Darfur_Sudan2, Kashmir1, Thailand1, Iran1, and Iraq1. Pairs of samples that can be modeled as independent gene pools were defined be a p-value ≥ 0.05. A second p-value threshold of between 0.01 and 0.05 was also taken in consideration as a possible statistically significant result.

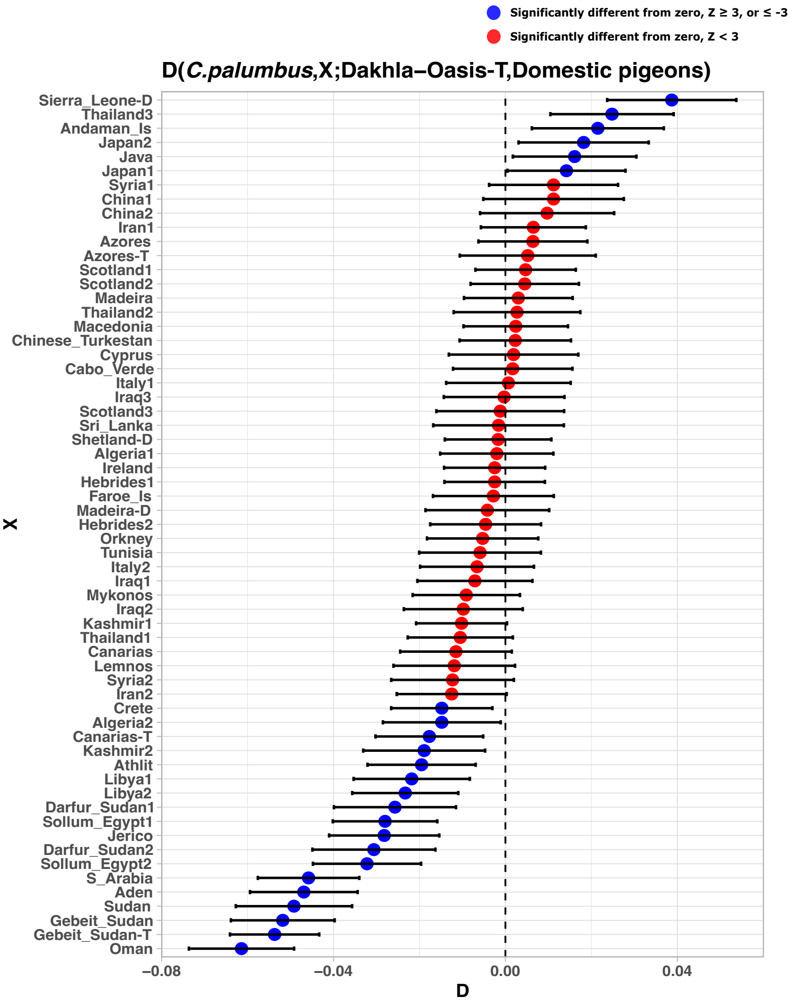


**Figure S7. Hybrid origin of *Columba livia dakhlae* type specimen.**
*D-statistics* analysis performed to test the close relationship observed between Dakhla-Oasis-T type specimen and the domestic breeds. The form of test was D(*C. palumbus*, X; Dakhlae-Oasis-T, Domestic pigeons), where *C. palumbus* was used as an outgroup, X represent all historical rock doves in our dataset and the domestic breeds that were grouped in a single population. Deviation from 0 was considered statistically significant when Z-score was below -3 or above 3. The significance of the test was assessed using a weighted block jackknife procedure over 1 Mb blocks.

Table S1. Samples metadata including the collection's information of each specimen, and the description of the processed genomic data.

| Project ID | Museum of origin | Museum ID | Species | Subspecies | Date | Sex | Locality | Tissue | Comment | Total number of pair reads | Total number of single-ended reads | Total number of hits | Total numbers of hits excluding PCR duplicates | Average read length per hit | Average genome coverage | DNA damage (%) at 3' G>A | DNA damage (%). 5' C>T |
|---|---|---|---|---|---|---|---|---|---|---|---|---|---|---|---|---|---|
| Iran1 | NHM Tring | 1934.1.1.1790 | *Columba livia* | *gaddi* | 1902 | Male | Fars, Iran | Dried Toepad | | 41206849 | 318119486 | 86041562 | 73151441 | 76.7543899 | 5.24834 | 2.2 | 2 |
| Algeria1 | NHM Tring | 1945:31:18 | *Columba livia* | *livia* | 1913 | female | El Kantara, Algeria | Dried Toepad | | 106436546 | / | 93954869 | 89636126 | 81.87541165 | 6.90442 | 1.5 | 1.5 |
| Sri_Lanka | NHM Tring | 1949:37:03 | *Columba livia* | *intermedia* | 1949 | Male | Kalametiya lagoon, Sri Lanka | Dried Toepad | | 37633724 | / | 30213076 | 27128664 | 65.81861145 | 1.45366 | 1.4 | 1.4 |
| Italy1 | NHM Tring | 1934.1.1.1794 | *Columba livia* | *livia* | 1920 | Male | San Severo di Puglia, Italy | Dried Toepad | | 43487737 | / | 35811754 | 32399678 | 60.05677414 | 1.66732 | 1.8 | 1.9 |
| Kashmir2 | NHM Tring | 1889.2.2.398 | *Columba livia* | *neglecta* | 1870 | Unknown | Kashmir | Dried Toepad | year of collection registration | 42944837 | / | 34316806 | 30721629 | 61.05288777 | 1.6238 | 3.7 | 3.6 |
| Hebrides1 | NHM Tring | 1965.M.4563 | *Columba livia* | *livia* | 1932 | Male | Grogarry, S. Uist, Hebrides | Dried Toepad | | 73495005 | / | 63505793 | 59416885 | 70.25388087 | 3.7448 | 1.4 | 1.4 |
| Mykonos | NHM Tring | 1934.4.17.22 | *Columba livia* | *livia* | 1933 | Male | Mykonos, Greece | Dried Toepad | | 44238513 | / | 36547875 | 33562660 | 61.4885271 | 1.84985 | 1.8 | 1.8 |
| Scotland1 | NHM Tring | 1897.11.6.38 | *Columba livia* | *livia* | 1897 | Male | Cromarty, Scotland | Dried Toepad | | 32482825 | / | 35711554 | 31526195 | 54.7145529 | 1.556031397 | 2.8 | 2.7 |
| Libya1 | NHM Tring | 1952.51.70 | *Columba livia* | / | 1952 | Unknown | Cyrene, Cyrenaica, Libya | Dried Toepad | | 78011947 | / | 27817458 | 25893521 | 62.00295788 | 1.45356 | 2.3 | 2.1 |
| Sollum_Egypt1 | NHM Tring | 1965.M.4568 | *Columba livia* | / | 1920 | Male | Sollum, West Egypty | Dried Toepad | | 68933491 | / | 66377007 | 61729921 | 63.91684244 | 3.63235 | 2.5 | 2.4 |
| Azores | NHM Tring | 1889.2.12.71 | *Columba livia* | / | 1865 | Unknown | St. Michaels, Azores | Dried Toepad | | 66149257 | / | 55947745 | 49076936 | 53.7930811 | 2.38305 | 2.3 | 2.2 |
| Cabo_Verde | NHM Tring | 1874.1.16.2 | *Columba livia* | / | 1874 | Unknown | Cabo Verde | Dried Toepad | year of collection registration | 80412558 | / | 56507721 | 51695688 | 60.79989974 | 2.82932 | 2.1 | 2.1 |
| Madeira | NHM Tring | 1965.M.4566 | *Columba livia* | / | 1924 | Male | S. Antonio, Madeira | Dried Toepad | | 73105903 | / | 68072564 | 63362674 | 72.78148399 | 4.2826 | 1 | 1.1 |
| Gebeit_Sudan | NHM Tring | 1915.12.24.254 | *Columba livia* | *schimperi* | 1912 | Male | Gebeit, Sudan | Dried Toepad | | 152336040 | / | 60307555 | 54756523 | 58.15214739 | 2.84193 | 1.6 | 1.6 |
| Canarias | NHM Tring | 1912.7.30.137 | *Columba livia* | / | 1909 | famale | (Tirajana) Gran Canarias | Dried Toepad | | 49608388 | / | 127151028 | 117932004 | 70.39426964 | 7.89682 | 2 | 2 |
| Athlit | NHM Tring | 1965-M-4572 | *Columba livia* | *gaddi* | 1920 | female | Athlit, Palestine | Dried Toepad | | 92249231 | / | 41536674 | 37612444 | 65.16238014 | 2.15011 | 1.2 | 1.3 |
| Iran2 | NHM Tring | 1965-M-4590 | *Columba livia* | *gaddi* | 1927 | Male | Birjand, Iran | Dried Toepad | | 152577852 | / | 77667173 | 72312522 | 67.4759045 | 4.6941 | 1.3 | 1.3 |
| Iraq1 | NHM Tring | 1927.4.16.2 | *Columba livia* | *gaddi* | 1927 | male | Baghdad, Iraq | Dried Toepad | | 70117302 | / | 129827812 | 88244095 | 69.37094156 | 7.57048 | 1.3 | 1.3 |
| Jerico | NHM Tring | 1965-M-4571 | *Columba livia* | *gaddi* | 1919 | Male | Jericho, Palestine | Dried Toepad | | 82466704 | / | 62663382 | 57933077 | 60.46152942 | 3.14747 | 2.7 | 2.6 |
| iraq3 | NHM Tring | 1965-M-4575 | *Columba livia* | *gaddi* | 1922 | Male | Dohuk, S. Kurdestan | Dried Toepad | | 133130197 | / | 69519398 | 62683371 | 60.29361677 | 3.43691 | 1.3 | 1.3 |
| Syria1 | NHM Tring | 1965-M-4570 | *Columba livia* | *gaddi* | 1919 | famale | Syrian desert. 40 miles E of Damascus | Dried Toepad | | 76470288 | / | 108737580 | 96307542 | 54.38848218 | 4.7573 | 1.3 | 1.3 |
| Tunisia | NHM Tring | 1925.12.25.8 | *Columba livia* | *gaddi* | 1925 | female | El Guettara, S.W. Tunisia | Dried Toepad | | 34172226 | / | 65747034 | 60152133 | 68.78355134 | 3.73506 | 1.1 | 1.1 |

| | | | | | | | | | | | | | | | | | |
|---|---|---|---|---|---|---|---|---|---|---|---|---|---|---|---|---|---|
| Libya2 | NHM Tring | 1952.51.67 | *Columba livia* | *gaddi* | 1952 | female | Slonta-tanamlu road, Cyrenaica, Libya | Dried Toepad | | 77181231 | / | 29530236 | 27798343 | 66.52181735 | 1.64367 | 1 | 1.3 |
| Siam1 | NHM Tring | 1955.1.1316 | *Columba livia* | *intermedia* | 1917 | Male | N' Chainat C. Siam, Thailand | Dried Toepad | | 71826882 | / | 62202265 | 52735848 | 55.70385711 | 2.64984 | 1.9 | 1.9 |
| Siam2 | NHM Tring | 1955.1.1312 | *Columba livia* | *intermedia* | 1919 | female | Bangkok, C. Siam, Thailand | Dried Toepad | | 66389729 | / | 53773937 | 47815246 | 56.13158354 | 2.3198 | 1.9 | 1.9 |
| Siam3 | NHM Tring | 1916.12.27.74 | *Columba livia* | *intermedia* | 1915 | female | Pak Jong, Siam, Thailand | Dried Toepad | | 32100110 | / | 52257591 | 46840181 | 63.15049245 | 2.53817 | 1.5 | 1.4 |
| Algeria2 | NHM Tring | 1923.8.7.1 | *Columba livia* | *livia* | 1913 | male | Jaya?, Algeria | Dried Toepad | specific location not found | 49246242 | / | 27424436 | 24905748 | 61.05979902 | 1.32118 | 1.1 | 1.1 |
| Faroe_Is | NHM Tring | 1897.10.20.2 | *Columba livia* | *livia* | 1878 | female | Faroe Islands | Dried Toepad | | 47167160 | / | 39325676 | 35399337 | 50.60348284 | 1.61214 | 2.3 | 2.2 |
| Hebrides2 | NHM Tring | 1953.76.100 | *Columba livia* | *livia* | 1931 | Male | Benbecula, O. Hebrides | Dried Toepad | | 39740654 | / | 40303043 | 36294858 | 57.57175001 | 1.8001 | 1.4 | 1.4 |
| Ireland | NHM Tring | 1934.1.1.1799 | *Columba livia* | *livia* | 1899 | Male | Co. Mayo, Ireland | Dried Toepad | | 34755498 | / | 32814777 | 30026577 | 63.7656425 | 1.71761 | 1.4 | 1.3 |
| Lemnos | NHM Tring | 1934.1.1.1793 | *Columba livia* | *livia* | 1907 | Male | Lemnos, Greece | Dried Toepad | | 62561518 | / | 29734459 | 25373019 | 54.16649985 | 1.11537 | 1.5 | 1.5 |
| Orkney | NHM Tring | 1881.5.1.5062 | *Columba livia* | *livia* | 1881 | Unknown | Orkney | Dried Toepad | year of collection registration | 19180366 | / | 49500000 | 45211074 | 69.96685986 | 2.74103 | 1.9 | 1.9 |
| Scotland2 | NHM Tring | 1934.9.1.1 | *Columba livia* | *livia* | 1937 | Male | Caithness, Scotland | Dried Toepad | | 51871016 | / | 16980653 | 16061078 | 69.82736837 | 1.011682667 | 1.1 | 1 |
| Italy2 | NHM Tring | 1965-M.4564 | *Columba livia* | *livia* | 1918 | female | Taranto, Italy | Dried Toepad | | 33399504 | 22431593 | 46078918 | 40189818 | 75.64101673 | 2.07794 | 1.5 | 1.4 |
| Macedonia | NHM Tring | 1934.11.20.211 | *Columba livia* | *livia* | 1933 | Male | Bistra Mountain, West of Macedonia | Dried Toepad | | 44023473 | 30970224 | 30755276 | 27560176 | 74.50748912 | 1.40234 | 1.4 | 1.3 |
| Crete | NHM Tring | 1965-M4567 | *Columba livia* | *livia* | 1920 | Male | Mount Ida, Crete | Dried Toepad | | 42969322 | 29139749 | 37725696 | 32412960 | 71.64327695 | 1.4515 | 1.7 | 1.6 |
| Ghana | NHM Tring | 1936.2.21.356 | *Columba livia* | *gymnocycla* | 1901 | Male | Gold Cost (Hinterland) | Dried Toepad | | 35756773 | 53772371 | 45534657 | 41723850 | 80.23745446 | 2.66716 | 1.7 | 1.7 |
| Mali | NHM Tring | 1932.8.6.36 | *Columba livia* | *gymnocycla* | 1932 | Male | Fiko, 30 miles East of Mopti, French Sudan | Dried Toepad | | 41208107 | 55161366 | 41547442 | 38152429 | 78.52274697 | 2.40762 | 1.4 | 1.4 |
| Kashmir1 | NHM Tring | 1949.Whi.1.1857 | *Columba livia* | *neglecta* | 1913 | female | Kashmir | Dried Toepad | | 35251782 | 40223745 | 44941744 | 39547246 | 82.3424964 | 2.64795 | 0.8 | 0.8 |
| Chinese_Turkestan | NHM Tring | 1931.7.8.12 | *Columba livia* | *neglecta* | 1930 | Male | Tekkes Valley, Chinese Turkestan | Dried Toepad | | 42463409 | 52427287 | 40527422 | 36723644 | 79.57089564 | 2.34588 | 1.5 | 1.6 |
| Shetland-D | NHM Tring | 1867.11.28.45 | *Columba livia* | / | 1867 | Unknown | Shetland | Dried Toepad | Darwin Collection | 72330929 | / | 36806678 | 33451860 | 52.71439715 | 1.43535 | 3.2 | 2.9 |
| Sierra_Leone-D | NHM Tring | 1867.11.28.48 | *Columba livia* | feral | 1867 | Unknown | Sierra Leone | Dried Toepad | Darwin Collection | 42214500 | / | 58149580 | 46263281 | 42.21378091 | 1.75896 | 3.5 | 3.4 |
| Madeira-D | NHM Tring | 1867.11.28.50 | *Columba livia* | feral | 1867 | Unknown | Madeira | Dried Toepad | Darwin Collection | 50622523 | / | 35356757 | 32944464 | 65.83022377 | 1.82864 | 2.4 | 2.2 |
| Japan1 | NHM Tring | 1884.1.16.3 | *Columba livia* | feral | 1880 | famale | Nagasaki, Japan | Dried Toepad | | 52877663 | / | 43025719 | 39889593 | 57.27883955 | 2.00008 | 2.5 | 2.3 |
| Java | NHM Tring | 1889.2.10.205 | *Columba livia* | feral | 1889 | Unknown | Java Island | Dried Toepad | year of collection registration | 43824586 | / | 40130530 | 34456275 | 52.27336114 | 1.56569 | 3 | 3 |
| Andaman_Is | NHM Tring | 1889.2.2.410 | *Columba livia* | feral | 1874 | Male | Port Blai, Andaman Islands | Dried Toepad | | 39835178 | / | 36338146 | 33392102 | 56.03803222 | 1.6237 | 2.5 | 2.4 |
| Japan2 | NHM Tring | 1897.10.30.449 | *Columba livia* | feral | 1897 | Unknown | Yokohama, Japan | Dried Toepad | | 49501509 | / | 34356035 | 30975851 | 53.6927552 | 1.500316481 | 2 | 2 |

| | | | | | | | | | | | | | | | | | |
|---|---|---|---|---|---|---|---|---|---|---|---|---|---|---|---|---|---|
| Azores-T | NHM Tring | 1904.12.31.308 | *Columba livia* | *atlantis* | 1903 | Male | Above Rosario, Corvo, Azores | Dried Toepad | Type specimen | 50787709 | / | 42252860 | 36894824 | 48.45770111 | 1.612769098 | 2.1 | 2 |
| China1 | NHM Tring | 1907.12.17.111 | *Columba livia* | *feral* | 1907 | Male | West of Wei-Hai-Wei, Shan-tung, China | Dried Toepad | | 63160437 | / | 43473718 | 40004354 | 61.01416048 | 2.201820167 | 2 | 1.8 |
| China2 | NHM Tring | 1908.1.2.56 | *Columba livia* | *feral* | 1898 | male | Shantung, China | Dried Toepad | | 39159388 | / | 54053741 | 47249202 | 49.9690043 | 2.129801444 | 2.2 | 2 |
| Cyprus | NHM Tring | 1909.8.7.1 | *Columba livia* | / | 1909 | Male | Athalassa, Cyprus | Dried Toepad | | 49811415 | / | 33704231 | 31215211 | 61.3414086 | 1.68341 | 1.7 | 1.6 |
| Canarias-T | NHM Tring | 1912.7.30.138 | *Columba livia* | *canariensis* | 1910 | female | Cueva de Las Ninas, Pinar Pajonal, Gran Canaria | Dried Toepad | Type specimen | 78952213 | / | 42126081 | 38049106 | 53.7301662 | 1.83933 | 2 | 1.9 |
| Gebeit_Sudan-T | NHM Tring | 1915.12.24.255 | *Columba livia* | *butleri* | 1912 | male | Gebeit, Red Sea Province, Sudan | Dried Toepad | Type specimen | 30658989 | / | 68160239 | 62229374 | 52.62233702 | 2.95078 | 2.2 | 2.2 |
| Sudan | NHM Tring | 1919.12.17.67 | *Columba livia* | / | 1914 | male | Sinkat, Red Sea Province, Sudan | Dried Toepad | | 34919589 | / | 26245518 | 24654681 | 58.19560298 | 1.28812 | 1.9 | 1.8 |
| Darfur_Sudan1 | NHM Tring | 1922.12.8.76 | *Columba livia* | *targia* | 1921 | male | Kurra, J. Marra, Darfur, Sudan | Dried Toepad | | 52390827 | / | 28868553 | 25118225 | 50.18055388 | 1.11694 | 1.7 | 1.8 |
| Mali-T | NHM Tring | 1932.8.6.1 | *Columba livia* | *lividor* | 1932 | male | Fiko, 30 miles East of Mopti, French Sudan | Dried Toepad | Type specimen | 43870264 | / | 45424064 | 41896689 | 60.20444584 | 2.25644 | 1.7 | 1.7 |
| Iraq2 | NHM Tring | 1933.2.16.216 | *Columba livia* | *palestinae* | 1918 | male | Mandali, Persian border (Iraq, Mesopotamia) | Dried Toepad | | 39287701 | / | 37688015 | 35095708 | 58.10737507 | 1.83115 | 1.9 | 1.9 |
| Syria2 | NHM Tring | 1939.12.9.812 | *Columba livia* | / | 1931 | female | Kahle m., 'Syrian desert' | Dried Toepad | | 24446091 | / | 34387756 | 31007365 | 51.40191011 | 1.41719 | 1.5 | 1.4 |
| Darfur_Sudan2 | NHM Tring | 1947-26-1 | *Columba livia* | *targia* | 1927 | male | Umdona, Darfur, Sudan | Dried Toepad | location not found | 36684606 | / | 20895728 | 18952420 | 51.24805024 | 0.815965 | 2.4 | 2.2 |
| Aden | NHM Tring | 1965-M-4592 | *Columba livia* | *palestinae* | 1922 | female | Aden | Dried Toepad | | 133961258 | / | 31231663 | 28816720 | 66.4150704 | 1.55454 | 1.8 | 1.7 |
| S_Arabia | NHM Tring | 1965-M-4601 | *Columba livia* | *palestinae* | 1948 | male | Ashaira, Western Saudi Arabia | Dried Toepad | | 32703830 | / | 114800182 | 106542935 | 78.34192053 | 8.17008 | 1.4 | 1.4 |
| Sollum_Egypt2 | NHM Tring | 1965-M.4569 | *Columba livia* | / | 1920 | Male | Sollum, West Egypty | Dried Toepad | | 43522073 | / | 28257098 | 26087632 | 61.0144988 | 1.30504 | 1.7 | 1.6 |
| Dakhla_Oasis-T | NHM Tring | 1965.M.10 | *Columba livia* | *dakhlae* | 1928 | female | Dakhla Oasis, Lybian Desert | Dried Toepad | Type specimen | 72348933 | / | 38244818 | 35696106 | 63.24226651 | 2.04703 | 1.4 | 1.5 |
| Scotland3 | NHM Tring | 2010.11.12 | *Columba livia* | / | 1913 | female | Ballantrea, Downau Point, Ayrshire, Scotland | Dried Toepad | | 31540531 | / | 61061456 | 55625905 | 57.5277518 | 2.95222 | 2 | 2.1 |
| Oman | NHM Tring | A/1998.92.15 | *Columba livia* | / | 1986 | Unknown | Jabal Qara', (Qaftaat)) Zufar, Oman | Toepad (alcohol) | | 42589351 | / | 30766534 | 28723278 | 67.13879854 | 2.0479 | 1.6 | 1.7 |

Table S2. Whole samples dataset included in this study.

| Project ID | Species | Location | Coverage | Sample | SAMN/SRA or Project numbers | Publication/Source |
|---|---|---|---|---|---|---|
| Iran1 | *Columba livia* | Fars, Iran | 5.24834 | mRD_01 | / | this study |
| Algeria1 | *Columba livia* | El Kantara, Algeria | 6.90442 | mRD_02 | / | this study |
| Sri_Lanka | *Columba livia* | Kalametiya lagoon, Sri Lanka | 1.45366 | mRD_03 | / | this study |
| Italy1 | *Columba livia* | Sancevro, Italy | 1.66732 | mRD_04 | / | this study |
| Kashmir2 | *Columba livia* | Kashmir, Pakistan | 1.6238 | mRD_05 | / | this study |
| Hebrides1 | *Columba livia* | Hebrides | 3.7448 | mRD_06 | / | this study |
| Mykonos | *Columba livia* | Mykonos, Greece | 1.84985 | mRD_07 | / | this study |
| Scotland1 | *Columba livia* | Scotland | 1.556031397 | mRD_08 | / | this study |
| Libya1 | *Columba livia* | Cyrene, Libya | 1.45356 | mRD_09 | / | this study |
| Sollum_Egypt1 | *Columba livia* | Sollum, Egypty | 3.63235 | mRD_10 | / | this study |
| Azores | *Columba livia* | Azores | 2.38305 | mRD_11 | / | this study |
| Syria1 | *Columba livia* | Syria | 2.82932 | mRD_12 | / | this study |
| Madeira | *Columba livia* | Madeira | 4.2826 | mRD_13 | / | this study |
| Gebeit_Sudan | *Columba livia* | Sudan | 2.84193 | mRD_14 | / | this study |
| Canarias | *Columba livia* | Canarias | 7.89682 | mRD_15 | / | this study |
| Athlit | *Columba livia* | Athlit, Palestine | 2.15011 | mRD_16 | / | this study |
| Iran2 | *Columba livia* | Birjand, Iran | 4.6941 | mRD_17 | / | this study |
| Iraq1 | *Columba livia* | Iraq | 7.57048 | mRD_18 | / | this study |
| Jerico | *Columba livia* | Jericho, Palestine | 3.14747 | mRD_19 | / | this study |
| Iraq3 | *Columba livia* | Kurdistan, Iraq | 3.43691 | mRD_20 | / | this study |
| Cabo_Verde | *Columba livia* | Cabo Verde | 4.7573 | mRD_21 | / | this study |
| Tunisia | *Columba livia* | Tunisia | 3.73506 | mRD_22 | / | this study |
| Libya2 | *Columba livia* | Cyrenaica, Libya | 1.64367 | mRD_23 | / | this study |
| Thailand1 | *Columba livia* | Thailand | 2.64984 | mRD_24 | / | this study |
| Thailand2 | *Columba livia* | Thailand | 2.3198 | mRD_25 | / | this study |
| Thailand3 | *Columba livia* | Thailand | 2.53817 | mRD_26 | / | this study |
| Algeria2 | *Columba livia* | Algeria | 1.32118 | mRD_27 | / | this study |
| Faroe_Is | *Columba livia* | Faroe Islands | 1.61214 | mRD_28 | / | this study |
| Hebrides2 | *Columba livia* | Hebrides | 1.8001 | mRD_29 | / | this study |
| Ireland | *Columba livia* | Ireland | 1.71761 | mRD_30 | / | this study |
| Lemnos | *Columba livia* | Lemnos, Greece | 1.11537 | mRD_31 | / | this study |
| Orkney | *Columba livia* | Orkney | 2.74103 | mRD_32 | / | this study |
| Scotland2 | *Columba livia* | Scotland | 1.011682667 | mRD_33 | / | this study |
| Italy2 | *Columba livia* | Taranto, Italy | 2.07794 | mRD_34 | / | this study |
| Macedonia | *Columba livia* | Macedonia | 1.40234 | mRD_35 | / | this study |
| Crete | *Columba livia* | Crete | 1.4515 | mRD_36 | / | this study |
| Ghana | *Columba livia* | Ghana | 2.66716 | mRD_37 | / | this study |
| Mali | *Columba livia* | Fiko, Mali | 2.40762 | mRD_38 | / | this study |
| Kashmir1 | *Columba livia* | Kashmir, Pakistan | 2.64795 | mRD_39 | / | this study |

| | | | | | | |
|---|---|---|---|---|---|---|
| Chinese_Turkestan | *Columba livia* | Turkestan, China | 2.34588 | mRD_40 | / | this study |
| Shetland-D | *Columba livia* | Shetland | 1.43535 | mRD_41 | / | this study |
| Sierra_Leone-D | *Columba livia* | Sierra Leone | 1.75896 | mRD_42 | / | this study |
| Madeira-D | *Columba livia* | Madeira | 1.82864 | mRD_43 | / | this study |
| Japan1 | *Columba livia* | Nagasaki, Japan | 2.00008 | mRD_44 | / | this study |
| Java | *Columba livia* | Java Island | 1.56569 | mRD_45 | / | this study |
| Andaman_Is | *Columba livia* | Andaman Islands | 1.6237 | mRD_46 | / | this study |
| Japan2 | *Columba livia* | Yokohama, Japan | 1.500316481 | mRD_47 | / | this study |
| Azores-T | *Columba livia* | Corvo, Azores | 1.612769098 | mRD_48 | / | this study |
| China1 | *Columba livia* | Shantung, China | 2.201820167 | mRD_49 | / | this study |
| China2 | *Columba livia* | Shantung, China | 2.129801444 | mRD_50 | / | this study |
| Cyprus | *Columba livia* | Cyprus | 1.68341 | mRD_51 | / | this study |
| Canarias-T | *Columba livia* | Pinar Pajonal, Gran Canaria | 1.83933 | mRD_52 | / | this study |
| Gebeit_Sudan-T | *Columba livia* | Gebeit, Sudan | 2.95078 | mRD_53 | / | this study |
| Sudan | *Columba livia* | Sinkat, Sudan | 1.28812 | mRD_54 | / | this study |
| Darfur_Sudan1 | *Columba livia* | Darfur, Sudan | 1.11694 | mRD_55 | / | this study |
| Mali-T | *Columba livia* | Fiko, Mali | 2.25644 | mRD_56 | / | this study |
| Iraq2 | *Columba livia* | Mandali, Iraq | 1.83115 | mRD_57 | / | this study |
| Syria2 | *Columba livia* | 'Syrian desert' | 1.41719 | mRD_58 | / | this study |
| Darfur_Sudan2 | *Columba livia* | Darfur, Sudan | 0.815965 | mRD_59 | / | this study |
| Aden | *Columba livia* | Aden, Yemen | 1.55454 | mRD_60 | / | this study |
| S_Arabia | *Columba livia* | Ashaira, Saudi Arabia | 8.17008 | mRD_61 | / | this study |
| Egypt_Sollum2 | *Columba livia* | Sollum, Egypty | 1.30504 | mRD_62 | / | this study |
| Dakhala_Oasis-T | *Columba livia* | Dakhla Oasis, Egypt | 2.04703 | mRD_63 | / | this study |
| Scotland3 | *Columba livia* | Ayrshire, Scotland | 2.95222 | mRD_64 | / | this study |
| Oman | *Columba livia* | Zufar, Oman | 2.0479 | mRD_65 | / | this study |
| Danish Tumbler | *Columba livia* | Denmark | 60.98 | BGI_PI-1 - 11 | BioSample: SAMN00993249; Sample name: BGI_PI-1; SRA: SRS333674 | Shapiro et al. 2013 |
| Indian fantail | *Columba livia* | / | 20.03290521 | BGI_idfan | BioSample: SAMN01057532; Sample name: BGI_idfan; SRA: SRS346864 | Shapiro et al. 2013 |
| Fantail | *Columba livia* | / | 23.01392705 | BGI_fan | BioSample: SAMN01057533; Sample name: BGI_fan; SRA: SRS346865 | Shapiro et al. 2013 |
| C. rupestris | *Columba rupestris* | / | 14.7358478 | BGI_C-rup | BioSample: SAMN01057534; Sample name: BGI_C-rup; SRA: SRS346866 | Shapiro et al. 2013 |
| Racing | *Columba livia* | / | 9.895150963 | BGI_PI-DI | BioSample: SAMN01057535; Sample name: BGI_PI-DI; SRA: SRS346867 | Shapiro et al. 2013 |
| Oriental | *Columba livia* | / | 6.774441222 | BGI_PI-DJ | BioSample: SAMN01057536; Sample name: BGI_PI-DJ; SRA: SRS346868 | Shapiro et al. 2013 |
| Laugher | *Columba livia* | / | 8.668028404 | BGI_PI-AC | BioSample: SAMN01057537; Sample name: BGI_PI-AC; SRA: SRS346869 | Shapiro et al. 2013 |
| Indian fantail | *Columba livia* | / | 8.554938798 | BGI_PI-AA | BioSample: SAMN01057538; Sample name: BGI_PI-AA; SRA: SRS346870 | Shapiro et al. 2013 |
| Feral (Salt Lake City, Utah, USA) | *Columba livia* | USA | 9.061812796 | BGI_PI-BC | BioSample: SAMN01057539; Sample name: BGI_PI-BC; SRA: SRS346871 | Shapiro et al. 2013 |
| English carrier | *Columba livia* | / | 10.39871739 | BGI_PI-AG | BioSample: SAMN01057540; Sample name: BGI_PI-AG; SRA: SRS346872 | Shapiro et al. 2013 |
| Scandaroon | *Columba livia* | / | 9.640421969 | BGI_PI-AH | BioSample: SAMN01057541; Sample name: BGI_PI-AH; SRA: SRS346873 | Shapiro et al. 2013 |
| Berlin longface tumbler | *Columba livia* | / | 10.13100487 | BGI_PI-AI | BioSample: SAMN01057542; Sample name: BGI_PI-AI; SRA: SRS346874 | Shapiro et al. 2013 |
| Birmingham roller | *Columba livia* | / | 8.905266226 | BGI_PI-AJ | BioSample: SAMN01057543; Sample name: BGI_PI-AJ; SRA: SRS346875 | Shapiro et al. 2013 |
| King | *Columba livia* | / | 10.81056783 | BGI_PI-AK | BioSample: SAMN01057544; Sample name: BGI_PI-AK; SRA: SRS346876 | Shapiro et al. 2013 |
| Chinese owl | *Columba livia* | / | 8.041128427 | BGI_PI-AL | BioSample: SAMN01057545; Sample name: BGI_PI-AL; SRA: SRS346877 | Shapiro et al. 2013 |
| Saxon monk | *Columba livia* | / | 11.91192554 | BGI_PI-AM | BioSample: SAMN01057546; Sample name: BGI_PI-AM; SRA: SRS346878 | Shapiro et al. 2013 |

| | | | | | | |
|---|---|---|---|---|---|---|
| Syrian dewlap | *Columba livia* | / | 11.1166817 | BGI_PI-AN | BioSample: SAMN01057547; Sample name: BGI_PI-AN; SRA: SRS346879 | Shapiro et al. 2013 |
| Shakhsharli | *Columba livia* | / | 11.46789002 | BGI_PI-AO | BioSample: SAMN01057548; Sample name: BGI_PI-AO; SRA: SRS346880 | Shapiro et al. 2013 |
| Oriental roller | *Columba livia* | / | 8.169535105 | BGI_PI-AP | BioSample: SAMN01057549; Sample name: BGI_PI-AP; SRA: SRS346881 | Shapiro et al. 2013 |
| Carneau | *Columba livia* | / | 10.59364749 | BGI_PI-AQ | BioSample: SAMN01057550; Sample name: BGI_PI-AQ; SRA: SRS346882 | Shapiro et al. 2013 |
| English longface tumbler | *Columba livia* | / | 7.668495123 | BGI_PI-AR | BioSample: SAMN01057551; Sample name: BGI_PI-AR; SRA: SRS346883 | Shapiro et al. 2013 |
| English pouter | *Columba livia* | / | 10.68001208 | BGI_PI-AS | BioSample: SAMN01057552; Sample name: BGI_PI-AS; SRA: SRS346884 | Shapiro et al. 2013 |
| Jacobin | *Columba livia* | / | 8.586987628 | BGI_PI-AT | BioSample: SAMN01057553; Sample name: BGI_PI-AT; SRA: SRS346885 | Shapiro et al. 2013 |
| Mookee | *Columba livia* | / | 10.69135875 | BGI_PI-AD | BioSample: SAMN01057554; Sample name: BGI_PI-AD; SRA: SRS346886 | Shapiro et al. 2013 |
| Starling | *Columba livia* | / | 9.533223161 | BGI_PI-AF | BioSample: SAMN01057555; Sample name: BGI_PI-AF; SRA: SRS346887 | Shapiro et al. 2013 |
| Spanish barb | *Columba livia* | / | 11.07154458 | BGI_PI-AE | BioSample: SAMN01057556; Sample name: BGI_PI-AE; SRA: SRS346888 | Shapiro et al. 2013 |
| African owl | *Columba livia* | / | 10.11713383 | BGI_PI-AB | BioSample: SAMN01057557; Sample name: BGI_PI-AB; SRA: SRS346889 | Shapiro et al. 2013 |
| ice pigeon | *Columba livia* | / | 11.62469438 | BGI_PI-BD | BioSample: SAMN01057558; Sample name: BGI_PI-BD; SRA: SRS346890 | Shapiro et al. 2013 |
| Frillback | *Columba livia* | / | 12.15460077 | BGI_PI-BE | BioSample: SAMN01057559; Sample name: BGI_PI-BE; SRA: SRS346891 | Shapiro et al. 2013 |
| iranian tumbler | *Columba livia* | / | 12.3772393 | BGI_PI-BF | BioSample: SAMN01057560; Sample name: BGI_PI-BF; SRA: SRS346892 | Shapiro et al. 2013 |
| Marchenero pouter | *Columba livia* | / | 13.80980787 | BGI_PI-BG | BioSample: SAMN01057561; Sample name: BGI_PI-BG; SRA: SRS346893 | Shapiro et al. 2013 |
| Runt | *Columba livia* | / | 13.61226882 | BGI_PI-BH | BioSample: SAMN01057562; Sample name: BGI_PI-BH; SRA: SRS346894 | Shapiro et al. 2013 |
| Saxon pouter | *Columba livia* | / | 13.64297974 | BGI_PI-BI | BioSample: SAMN01057563; Sample name: BGI_PI-BI; SRA: SRS346895 | Shapiro et al. 2013 |
| English trumpeter | *Columba livia* | / | 12.76043828 | BGI_PI-BJ | BioSample: SAMN01057564; Sample name: BGI_PI-BJ; SRA: SRS346896 | Shapiro et al. 2013 |
| Lahore | *Columba livia* | / | 11.92217449 | BGI_PI-AU | BioSample: SAMN01057565; Sample name: BGI_PI-AU; SRA: SRS346897 | Shapiro et al. 2013 |
| Lebanon | *Columba livia* | / | 10.93237512 | BGI_PI-AV | BioSample: SAMN01057566; Sample name: BGI_PI-AV; SRA: SRS346898 | Shapiro et al. 2013 |
| Parlor roller | *Columba livia* | / | 7.474224178 | BGI_PI-AW | BioSample: SAMN01057567; Sample name: BGI_PI-AW; SRA: SRS346899 | Shapiro et al. 2013 |
| Racing homer | *Columba livia* | / | 11.78342911 | BGI_PI-AX | BioSample: SAMN01057568; Sample name: BGI_PI-AX; SRA: SRS346900 | Shapiro et al. 2013 |
| Archangel | *Columba livia* | / | 11.78186737 | BGI_PI-AY | BioSample: SAMN01057569; Sample name: BGI_PI-AY; SRA: SRS346901 | Shapiro et al. 2013 |
| Cumulet | *Columba livia* | / | 9.906565827 | BGI_PI-AZ | BioSample: SAMN01057570; Sample name: BGI_PI-AZ; SRA: SRS346902 | Shapiro et al. 2013 |
| Egyptian swift | *Columba livia* | / | 10.44490396 | BGI_PI-BA | BioSample: SAMN01057571; Sample name: BGI_PI-BA; SRA: SRS346903 | Shapiro et al. 2013 |
| Feral (Lake Anna, Virginia, USA) | *Columba livia* | USA | 11.62122622 | BGI_PI-BB | BioSample: SAMN01057572; Sample name: BGI_PI-BB; SRA: SRS346904 | Shapiro et al. 2013 |
| C. guinea | *Columba guinea* | / | 21.88301951 | Cogui | BioSample: SAMN04886479; Sample name: Cogui; SRA: SRS1416880 | Vickrey et al. 2018 |
| C. larvata | *Columba larvata* | / | 3.52264353 | Aplar | BioSample: SAMN04886481; Sample name: Aplar; SRA: SRS1416882 | Vickrey et al. 2018 |
| C. palumbus | *Columba palumbus* | / | 20.12975157 | Copal | BioSample: SAMN04886480; Sample name: Copal; SRA: SRS1416881 | Vickrey et al. 2018 |
| P. fasciata | *Patagioenas fasciata* | USA | 3.747136425 | BTP2013 | BioSample: SAMN04386172; Sample name: BTP2013; SRA: SRS2103361 | Murray et al. 2017 |
| P. colchicus | *Phasianus colchicus* | China | 6.28662773 | Phasianus colchicus whole genome sequencing data | BioSample: SAMN08888528; Sample name: Phasianus colchicus whole genome sequencing data; SRA: SRP139710 | Liu et al. 2019 |
